# Evolution of median fin patterning and modularity in living and fossil osteichthyans

**DOI:** 10.1101/2022.07.18.500482

**Authors:** France Charest, Jorge Mondéjar Fernández, Thomas Grünbaum, Richard Cloutier

**Author notes:** Corresponding author (FC). These authors contributed equally to this work.

## Abstract

Morphological and developmental similarities, and interactions among developing structures are interpreted as evidences of modularity. Such similarities exist between the dorsal and anal fins of living actinopterygians: (1) both fins differentiate in the same direction [dorsal and anal fin patterning module (DAFPM)], and (2) radials and lepidotrichia differentiate in the same direction [endoskeleton and exoskeleton module (EEM)]. To infer the evolution of these common developmental patternings among osteichthyans, we address (1) the complete description and quantification of the DAFPM and EEM in a living actinopterygian (the rainbow trout *Oncorhynchus mykiss*) and (2) the presence of these modules in fossil osteichthyans (coelacanths, lungfishes, porolepiforms and ‘osteolepiforms’). In *Oncorhynchus*, sequences of skeletal elements are determined based on (1) apparition (radials and lepidotrichia), (2) chondrification (radials), (3) ossification (radials and lepidotrichia), and (4) segmentation plus bifurcation (lepidotrichia). Correlations are then explored between sequences. In fossil osteichthyans, sequences are determined based on (1) ossification (radials and lepidotrichia), (2) segmentation, and (3) bifurcation of lepidotrichia. Segmentation and bifurcation patterns were found crucial for comparisons between living and extinct taxa. Our data suggest that the EEM is plesiomorphic at least for actinopterygians, and the DAFPM is plesiomorphic for osteichthyans, with homoplastic dissociation. Finally, recurrent patterns suggest the presence of a Lepidotrichia Patterning Module (LPM).

## Introduction

In the past two decades, the median fins [i.e., dorsal, anal, and caudal fins] of fishes have been the focus of an overwhelming body of research in evolutionary developmental biology. Primary interest for these so-called unpaired fins lies in their locomotor functions [1–3], ecological implications [4, 5], comparative morpho-anatomy [6], molecular [7] and developmental [8] patterning, as well as morphological disparity [9]. Indeed, among median fins, the dorsal and anal fins of piscine osteichthyans show a great morphological disparity, reflecting the evolvability of this system [9–11].

Osteichthyans primitively display two dorsal fins and a single anal fin [12]. Independently and repeatedly in actinopterygians and sarcopterygians, the number of dorsal fins is reduced by loss or fusion with the caudal fin [9,11,13]. The loss of the anal fin is less frequent in piscine osteichthyans, although loss or fusion with the caudal fin occurs in some teleosts and dipnoans [11, 14]. Among ‘elpistostegalians’, the extinct transitional taxa between fishes and tetrapods, the condition is poorly documented (e.g., *Panderichthys, Tiktaalik*). However, *Elpistostege*, considered a basal tetrapod, has lost the dorsal fin while an anal fin is present [15]. And finally, the absence of both the dorsal and anal fins is considered a synapomorphy shared by aquatic and terrestrial tetrapods [14] with the exception of *Elpistostege* [15]. Even with such morphological disparity, the structure and development of the median fins is expected to be broadly similar among osteichthyans because these fins have similar constituents.

In a forerunner comparative study, Mabee et al. [16] revealed the recurrence of similar developmental patterning (i.e., sequences and direction of development among endoskeletal and exoskeletal elements) in the dorsal and anal fins among living actinopterygians, which they interpreted as evidence of modularity. They found out that the similar patterning of these fins might be indicative of two modules: (1) the Dorsal and Anal Fin Patterning Module (DAFPM), where the skeletal elements (radial bones and lepidotrichia) of both fins differentiate in the same direction and (2) the Endoskeleton and Exoskeleton Module (EEM), where the directions of development of the endoskeleton (radials) and exoskeleton (lepidotrichia) are similar. The DAFPM and EEM are considered to be maintained during actinopterygian phylogeny [16] but a phylogenetic inference across osteichthyans was not possible owing to the absence of comparative data on early actinopterygians and sarcopterygians. A broad phylogenetic sampling of actinopterygians and sarcopterygians is thus necessary to document the patterns of developmental similarity between dorsal and anal fins throughout osteichthyan evolution. However, in order to validate the prospective existence and distribution of median fin modules among osteichthyans (i.e., specifically the DAFPM and EEM *sensu* [16]), it is mandatory to include extant as well as extinct taxa within a comparative framework.

The rarity of fossilized ontogenies [17] and a bias toward the preservation of hard tissues (bones *versus* cartilages) limit our assessment of early developmental patterns in extinct taxa. However, the presence of fossilized individual ontogenies (i.e., anatomical structures found in adults that have recorded individual developmental patterns), can potentially broaden the phylogenetic sampling. The complex structure of osteichthyan fin rays (e.g., [18, 19]) provides such developmental data because fin rays are accretional structures for which structural elements are added without substantial remodelling, thus allowing the preservation of early developmental patterning.

The main objectives of this study are (1) to provide a complete description of the patterns of developmental similarity of the dorsal and anal fins in a living actinopterygian, the rainbow trout (*Oncorhynchus mykiss*), (2) to describe the dorsal and anal fins patterning in fossil osteichthyans, and (3) to compare the similarity of the developmental patterns in order to discuss the prospective existence of the DAFPM and EEM modules within osteichthyans. Patterns of developmental similarity were investigated by using relative developmental sequences and direction of development among endoskeletal and exoskeletal elements of the dorsal and anal fins. We expect developmental sequences to be (1) significantly congruent between dorsal and anal fins (i.e., indicative of DAFPM), and (2) significantly congruent between the endoskeleton and exoskeleton within each fin (i.e., indicative of EEM).

### Nomenclature

In order to facilitate comparisons during the description of fin morphologies and its constituents, a brief review of the main elements composing the median fins of osteichthyans is presented here. We will detail the different types of fins rays encountered in osteichthyans (i.e., lepidotrichia and actinotrichia) and the events used to define the median fin developmental patterning.

#### Lepidotrichia

Lepidotrichia are osseous fin rays of dermal origin. Each lepidotrichium is composed of two parallel and symmetrical elements called hemirays. Lepidotrichia are usually segmented (i.e., “jointed” lepidotrichia). The “joints” correspond to very narrow, non-mineralized spaces between adjacent segments connected by collagenous ligaments called Sharpey’s fibres. Adjoining segments are sequentially added distally during growth [20] before the ossification of the ray, which begins proximally. The most proximal segment is always longer than the others from the very first stages of development. This pattern has been observed in many living and fossil species (e.g., the actinopterygians *Gobius*, *Pygosleus*, *Cottus*, and *Blennius*, among others, and the sarcopterygian *Miguashaia* and *Eusthenopteron* [21–23]. The lepidotrichia articulate with the most distal endoskeletal elements (e.g., radials or pterygiophores; or the phalanges as in *Elpistostege* [15]) in the paired and median fins. The hemirays are contralaterally arranged on both sides of the endoskeleton. Usually, each radial bone carries more than one lepidotrichium, however certain derived taxa (e.g., teleosts and coelacanths) show a 1:1 ratio between the radials and the fin rays in the median fins. In their most distal portion, the lepidotrichia usually bifurcate (i.e., branched lepidotrichia) as if the ray was split in two. Numerous episodes (or orders) of bifurcation can occur in a single ray. Lepidotrichia represent a synapomorphy of crown osteichthyans [24]. Extant dipnoans (i.e., *Neoceratodus*, *Lepidosiren* and *Protopterus*) display a unique kind of partially ossified lepidotrichia termed camptotrichia [25].

#### Actinotrichia

Actinotrichia are flexible fin rays formed by long fibres of collagen known as elastoidine [19]. The actinotrichia form the main support of the osteichthyan fins in larval and juvenile stages of the ontogeny and are found in the most distal part of the adult fins arranged in contralateral palisades. During the formation of the lepidotrichia, actinotrichia are progressively resorbed, both within hemirays and between lepidotrichia, leaving only a narrow distal fringe. The formation of actinotrichia is followed by the apparition and development of the endoskeletal elements (e.g., radials), and the formation of the lepidotrichia [16,26–28]. Mesenchymal cells (osteoblasts) may then use the actinotrichia as a scaffold during the initial stages of formation of the lepidotrichia [19,26,29].

#### Median fin patterning

Median fin patterning can be defined in terms of a series of events rather than solely on formation (or differentiation, *sensu* [16]). An event defines a unit of transformation with concomitant phenotypic changes (e.g., lepidotrichia ossification) and an event may have different properties (e.g., onset, offset, duration) [30]. The chronological order of events corresponds to a sequence. Fins being composed of different elements, we referred to a developmental sequence when comparing the same developmental state (e.g., apparition, chondrification, ossification) among elements. We referred to an ontogenetic sequence when comparing different developmental states for a single element [31].

Of the numerous developmental events associated with fin formation in living actinopterygians [8,28,32–38], our study focused on 11 skeletogenic events: (1) apparition (i.e., collagenous matrix precursor) of actinotrichia, (2) apparition (i.e., mesenchymal condensation) of proximal radials, (3) chondrification of proximal radials, (4) apparition of lepidotrichia, (5) apparition (i.e., mesenchymal condensation) of distal radials, (6) chondrification of distal radials, (7) segmentation of lepidotrichia, (8) ossification of lepidotrichia, (9) bifurcation of lepidotrichia, (10) ossification of proximal radials, and (11) ossification of distal radials. The establishment of these developmental events is a powerful tool that could be used to uncover patterns of developmental similarity and shed light on patterns of modularity across osteichthyan evolution.

## Material and methods

### Living material

Developmental sequences of the dorsal and anal fins were obtained from embryo- juvenile specimens of the rainbow trout (*Oncorhynchus mykiss*) ranging from 5 days pre- hatching to 100 days post-hatching (dph). Alevins-juveniles were reared in swimming channels under constant water velocity (0.4 cm/s) in 2005 [see [39] for rearing conditions]. Specimens were sampled every day up to 34 dph, every other day from 34 to 80 dph, and every four days up to 100 dph. Samples were fixed in neutral buffered formalin for 48h, and then preserved in 70% ethanol. One specimen for each sampling day, plus one or two replicates for specimens between 0 to 24 dph, were cleared-and-double stained with Alizarin red S for bones and Alcian blue for cartilages [40]. Replicates were used to palliate with staining problems [41]. Pre-hatching specimens were removed from their egg capsule prior to clearing and Alcian blue staining. Digital pictures were taken before staining and 5- 10 days after staining to avoid interpretive errors owing to destaining. In total, eighty specimens were used to reconstruct developmental sequences of serial skeletal elements from the dorsal and anal fins (Figs 1 and 2) for all the events associated with fin development. All specimens were reared and used for a previous experiment [39], for which protocols were approved by the Université du Québec à Rimouski’s animal care and use committee.

**Fig 1.**
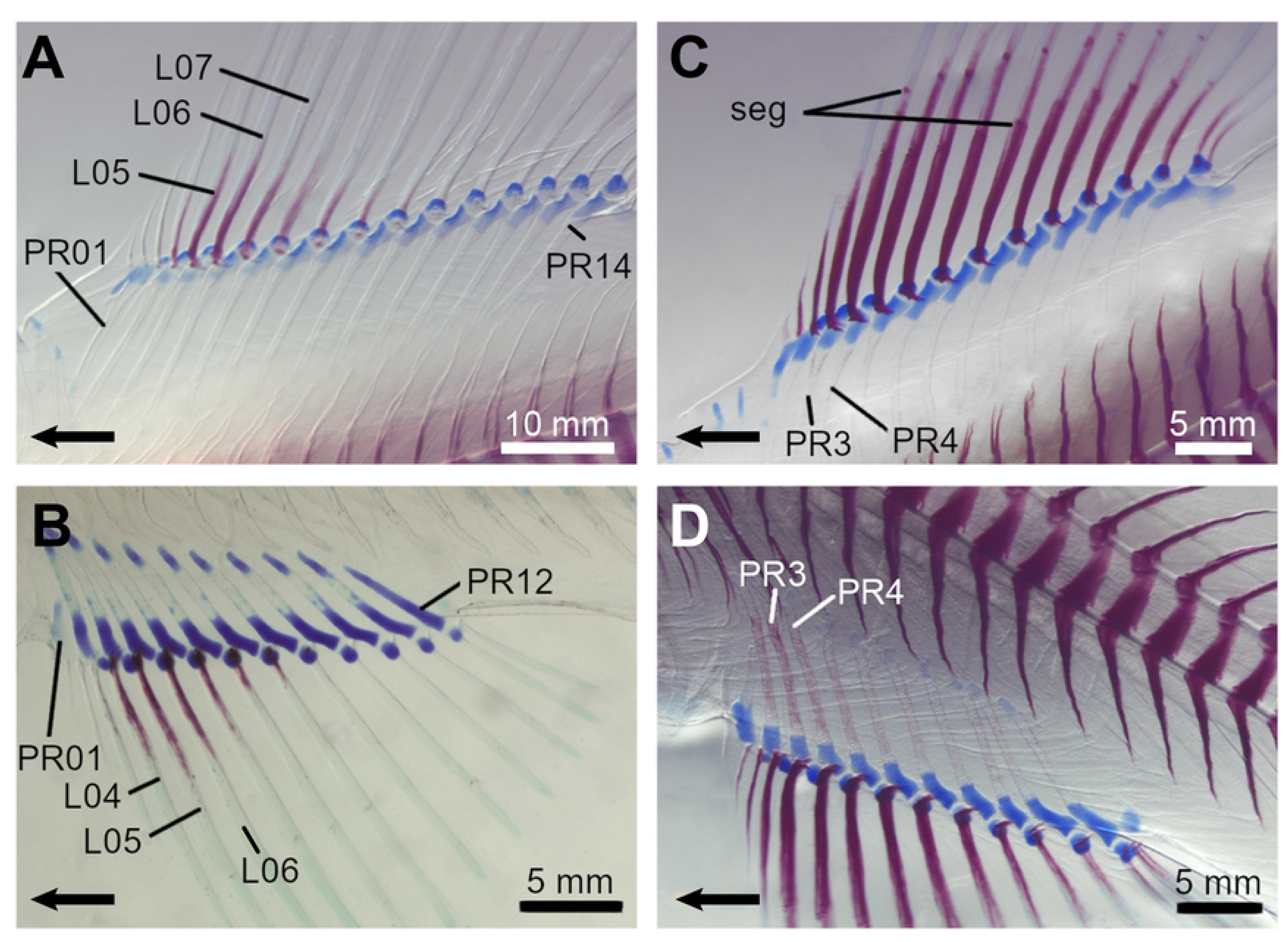
Details of the dorsal and anal fins of the rainbow trout (*Oncorhynchus mykiss*) based on a 31.33 mm long juvenile specimen. The main morphological features of these fins are identified. The serial elements, the proximal radials (PR), the distal radials (DR) and the lepidotrichia (L), are numbered from the anterior to the posterior of the fins. Act, actinotrichia; an.f, anal fin; bif, bifurcation; caud.f, caudal fin; DR, distal radial; dors.f, dorsal fin; L, lepidotrichia; PR, proximal radial; pect.f, pectoral fin; pelv.f, pelvic fin; seg, segmentation.

**Fig 2.**
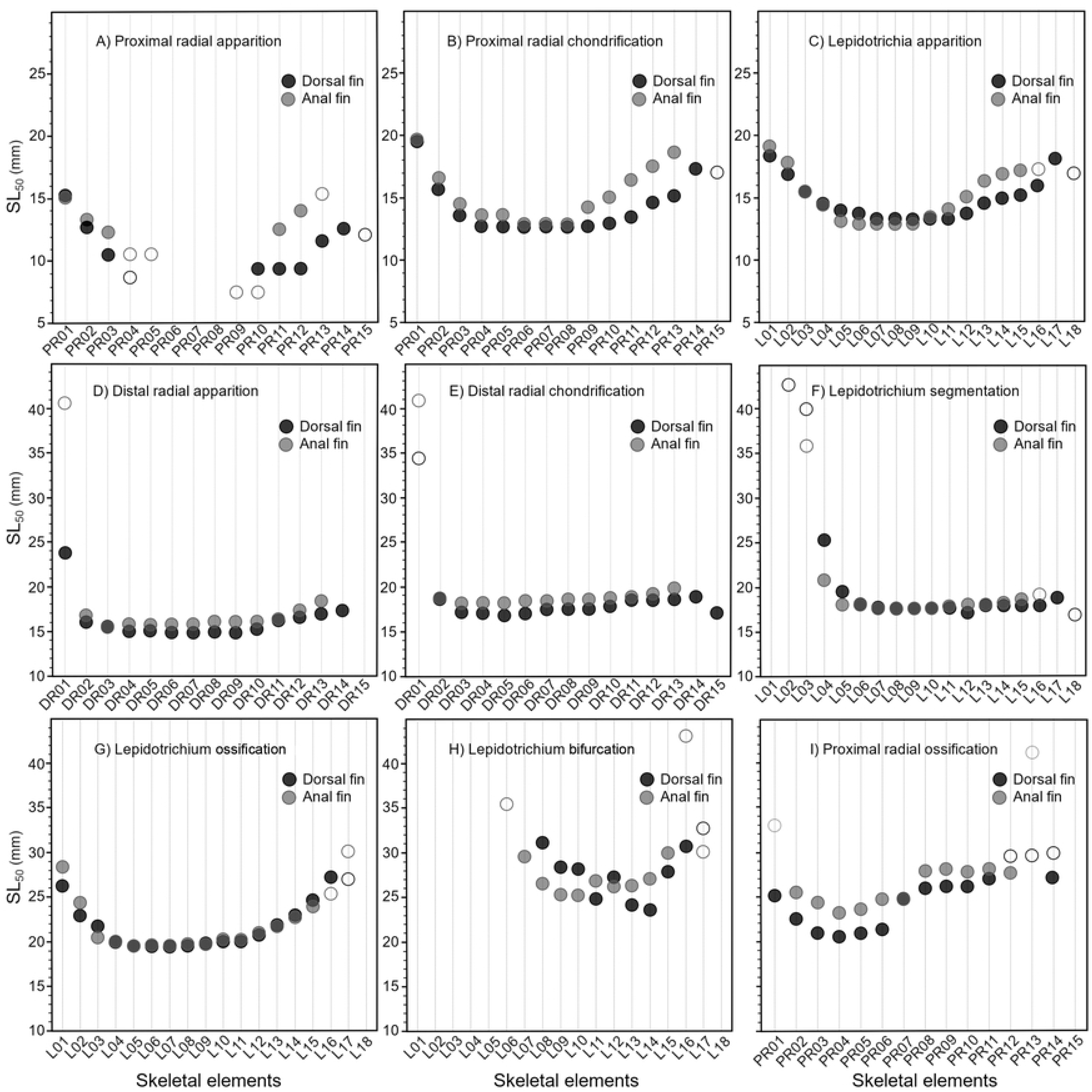
Cleared and stained specimens of the rainbow trout (*Oncorhynchus mykiss*). The cartilage is stained in red and the cartilages in blue. A: Dorsal fin showing the beginning of lepidotrichia ossification (specimen SL = 28.36 mm); B: Anal fin showing the beginning of lepidotrichia ossification (specimen SL = 19.80 mm); C: Dorsal fin showing the beginning of radial ossification (specimen SL = 24.90 mm); D: Anal fin showing the beginning of radial ossification (specimen SL = 24.90 mm). Arrows point anteriorly. L, lepidotrichia; PR, proximal radial; seg, segmentation.

Observations were made under a Leica MZ16A binocular mounted with a digital camera. Standard length (SL) was measured prior to staining with Northern Eclipse Software (Version 6.0). Since SL and dph are highly correlated (*r*^2^ = 0.952; *P* < 0.001) and SL is recognized as a better proxy for morphological development in fishes [42], SL was used for all statistical analyses.

Event coding was based on colour uptake by skeletal elements. In this study, three states are recognized for radials: (1) present (mesenchymal cell condensations without stain uptake), (2) cartilaginous (blue), and (3) ossified (red). Four developmental states are recognized for lepidotrichia: (1) present (collagenous matrix), (2) ossified (red), (3) segmented (number of segments per lepidotrichium), and (4) bifurcated (position of the bifurcation along the lepidotrichia). Surveyed specimens with their size (SL) and event coding of the skeletal elements are listed in S1 File.

To manage with the inter-individual variation in the number of radials and lepidotrichia, positional homologies and numbering of elements were inferred *a posteriori* by lining up all specimens with the third radial (variation being more important in peripheral areas) and by comparing similarities among sequences of similar-sized specimens. Myomere counts (from cranial to caudal) were used in the earliest stages as a topographical criterion to identify the first proximal radials to differentiate. The dorsal and anal fins are positioned at the level of myomeres 21-32 and 40-50, respectively.

Logistic regressions were used to estimate the SL at which 50% (SL_50_) of the specimens have reached a given developmental state (i.e., present, cartilaginous, ossified, segmented and bifurcated) for each skeletal element (see [30] for further details).

Significance of the logistic regressions were tested using the Likelihood ratio statistic [43]. To interpret a regression for a given element, the significance level was calculated using the Bonferroni correction; the collective significance level of 0.05 was divided by the number of elements to get the nominal significance level for each regression. Statistical analyses were performed with R Studio for Windows v. 1.3.1093 (library: MASS R; [44]).

In order to investigate the median fins patterning and developmental similarity, the SL_50_ values of each skeletal element (i.e., derived from the logistic regressions) were used to order the serial elements in relative developmental sequences within a fin. The relative order of a skeletal element within a developmental sequence was then converted by attributing a rank value. Spearman rank correlation coefficients were then used to describe the relations between the developmental sequences in the dorsal and anal fins, and in the endoskeleton and exoskeleton (see [30, 45] for the detailed procedure). Only the elements for which the logistic model was significant (under the nominal significance level) were considered for Spearman correlations. Logistic regressions do not produce SL_50_ when the elements are present in all specimens. The actual sizes of the smallest specimens were included in the Spearman correlations involving the apparition of the proximal radials in the dorsal and anal fins, since few skeletal elements were already present in these specimens. These already present proximal radials are ranked as the first appeared in the developmental sequence for the Spearman correlations.

### Fossil material

The phylogenetic sampling includes six Palaeozoic (Devonian-Carboniferous) species of osteichthyans comprising a ‘palaeonisciform’ actinopterygian (*Elonichthys peltigerus*) and five sarcopterygians including coelacanths (*Miguashaia bureaui* and *Rhabdoderma exiguum*), lungfishes (*Dipterus valenciennesi*), porolepiforms (*Quebecius quebecensis*) and ‘osteolepiforms’ (*Eusthenopteron foordi*). Specimens were chosen according to their exceptional state of preservation (articulated postcranial material and undistorted fins) and, whenever possible, availability of ontogenetic series. Specimens of *E. peltigerus* and *R. exiguum* come from the Upper Carboniferous (middle Pennsylvanian) Francis Creek Shale (Mazon Creek area, Illinois) and studied specimens are housed in the Field Museum of Natural History (FMNH; Chicago, IL, USA). Specimens of *M. bureaui*, *Q. quebecensis* and *E. foordi* come from the Upper Devonian (middle Frasnian) Escuminac Formation (Miguasha, Quebec, Canada); studied specimens are housed in the Musée d’Histoire Naturelle de Miguasha (MHNM, parc national de Miguasha, Quebec, Canada), the American Museum of Natural History (AMNH, New York, NY, USA), the Musée de géologie René-Bureau from Université Laval (ULQ, Quebec, Canada), and The University of Kansas Biodiversity Institute and Museum of Natural History, Division of Vertebrate Paleontology (KUVP, Lawrence, KS, USA). Finally, specimens of *D. valenciennesi* come from the Middle Devonian (Givetian) Achanarras beds (Scotland, UK); studied specimens are housed in the Natural History Museum (BMNH, London, UK). A complete list of surveyed specimens with their size [SL or total length (TL)] are listed in S1 Table.

Fossil specimens were examined under a Leica MZ9.5 binocular equipped with a drawing tube and were photographed with an Olympus Camedia C5060. Developmental states include (1) ossification (presence; radials and lepidotrichia), (2) segmentation (lepidotrichia), and (3) bifurcation (lepidotrichia).

## Results

### Developmental patterning of the dorsal and anal fins of

#### Oncorhynchus

The dorsal and anal fins of juvenile *Oncorhynchus* are similar in their anatomy and shape but differ slightly in meristic counts of skeletal elements and fin size (Fig 1; Table 1). Generally, the dorsal and anal fins are composed of 29 (i.e., 15 proximal and 14 distal radials) and 25 (i.e., 13 proximal and 12 distal radials) endoskeletal elements, and 18 and 15 exoskeletal elements (i.e., lepidotrichia), respectively. In both fins, proximal radials (PR), distal radials (DR), and lepidotrichia (L) are organized in a one-to-one relationship, with the exception of the first proximal and distal radials that support four and three lepidotrichia in the dorsal and anal fins, respectively, while the last proximal and distal radials support two lepidotrichia in both fins. Nine of the eleven developmental events previously described (see Nomenclature) were analysed in *Oncorhynchus*; the relative developmental sequence in each fin is ordered based on SL_50_ (Table 2). The presence of actinotrichia (event 1) was not analysed except for its initial position in the sequence of events. The ossification of distal radials (event 11) occurs after 100 dph, beyond the timeframe of our study thus no data were available.

**Table 1.** Meristic counts for the skeletal elements, proximal radials (PR), distal radials (DR) and lepidotrichia (L) and proportions of the dorsal (D) and anal (A) fins in living and extinct osteichthyans.

| Taxa | Dorsal fin |  |  | Anal fin |  |  | Proportion |
| --- | --- | --- | --- | --- | --- | --- | --- |
|  | PR | DR | L | PR | DR | L |  |
| *Oncorhynchus* | 12-15 | 12-14 | 15-18 | 11-13 | 11-13 | 12-16 | D > A |
| *Elonichthys* | 5-12 | 1-17 | 34-40 | 3-9 | 1-17 | 40-47 | D < A |
| *Miguashaia* | 1 | NA | 21-28 | 1 | NA | 19-25 | D = A |
| *Rhabdoderma* | ? | ? | 12-21 | ? | ? | 9-21 | D = A |
| *Quebecius* | ? | ? | 30-35 | ? | ? | 31-36 | D = A |
| *Dipterus* | 1<sup>1</sup> | 5<sup>1</sup> | 42-48 | 1<sup>1</sup> | 4<sup>1</sup> | 22-28 | D > A |
| *Eusthenopteron* | 1 | 3 | 19-26 | 1 | 3 | 20-25 | D = A |

NA, non applicable
?, endoskeletal elements are unknown
<sup>1</sup> Following Ahlberg and Trewin [46]; there is more than one row of distal radials; data are for one row.

**Table 2.**
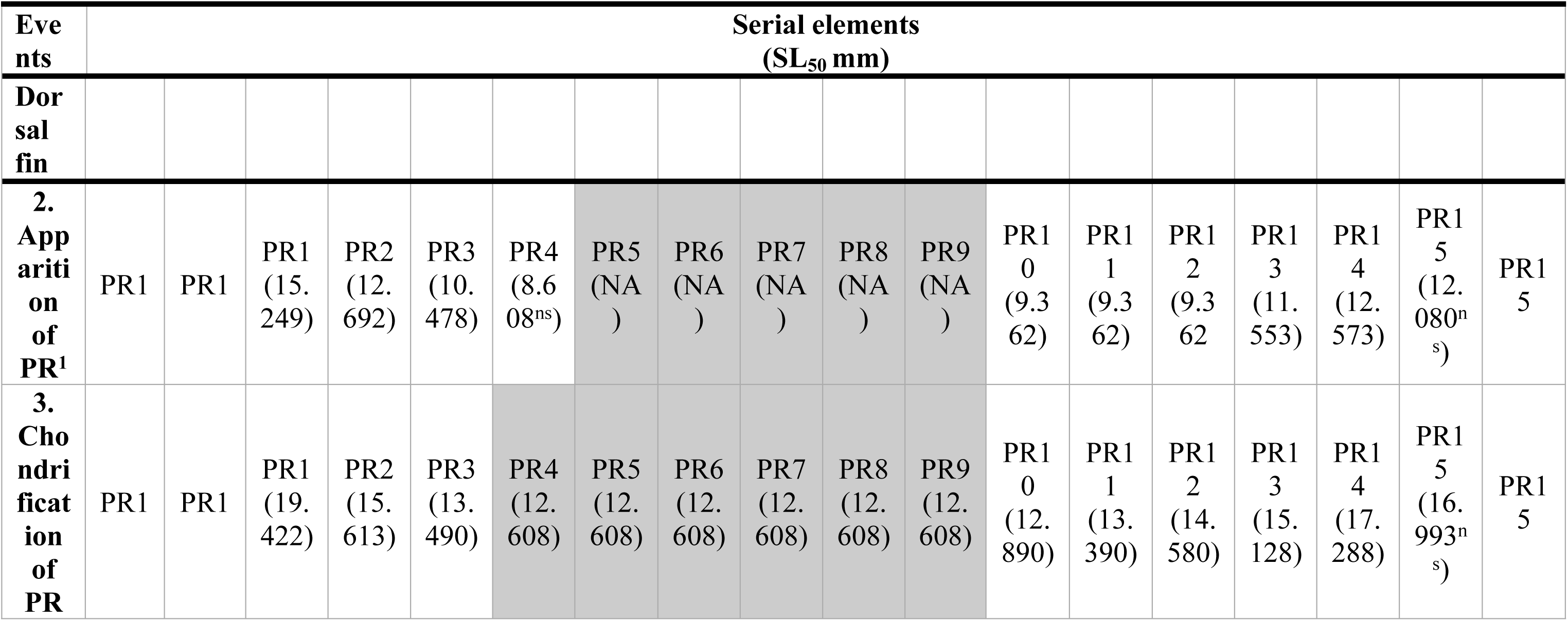
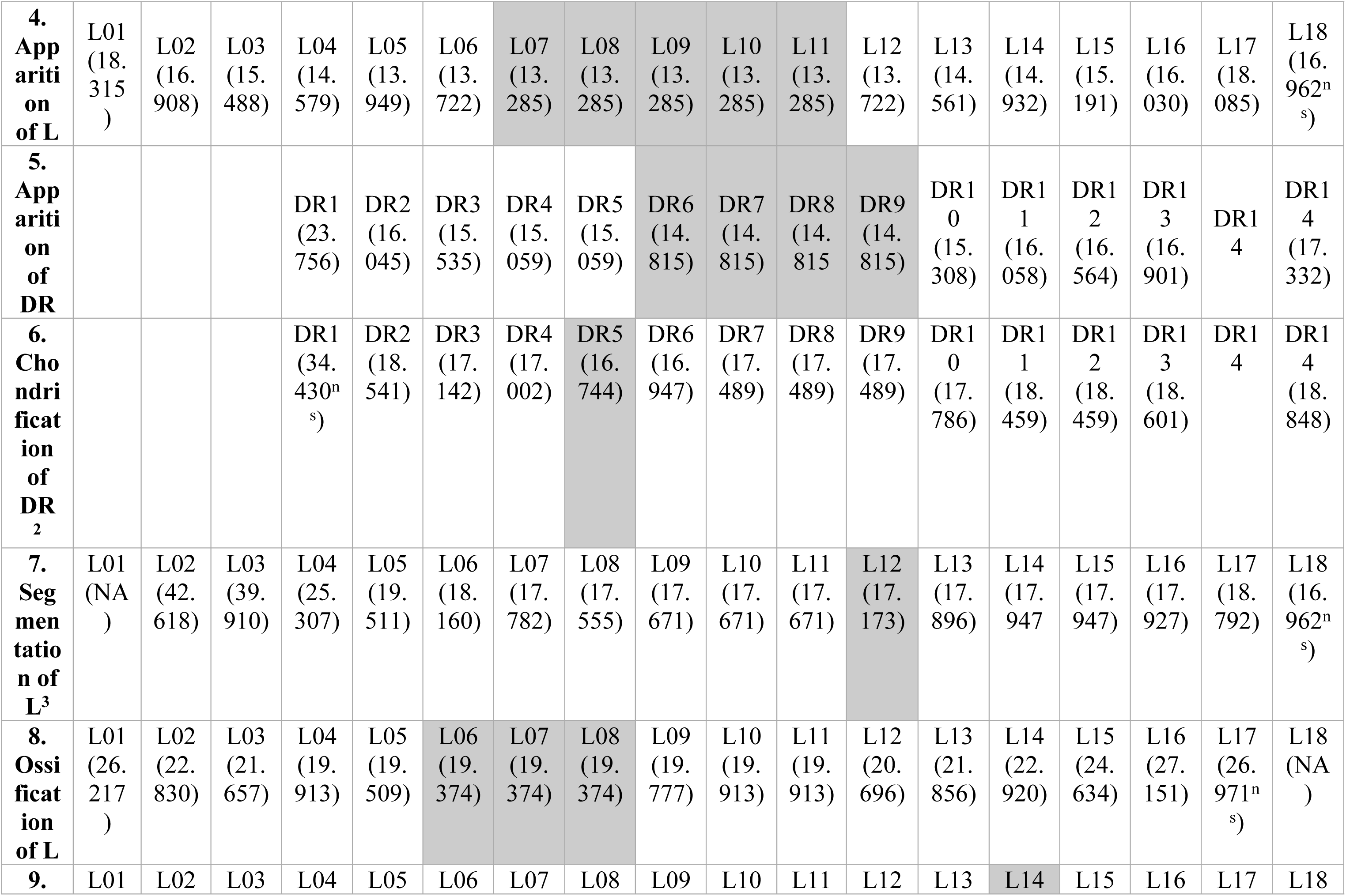
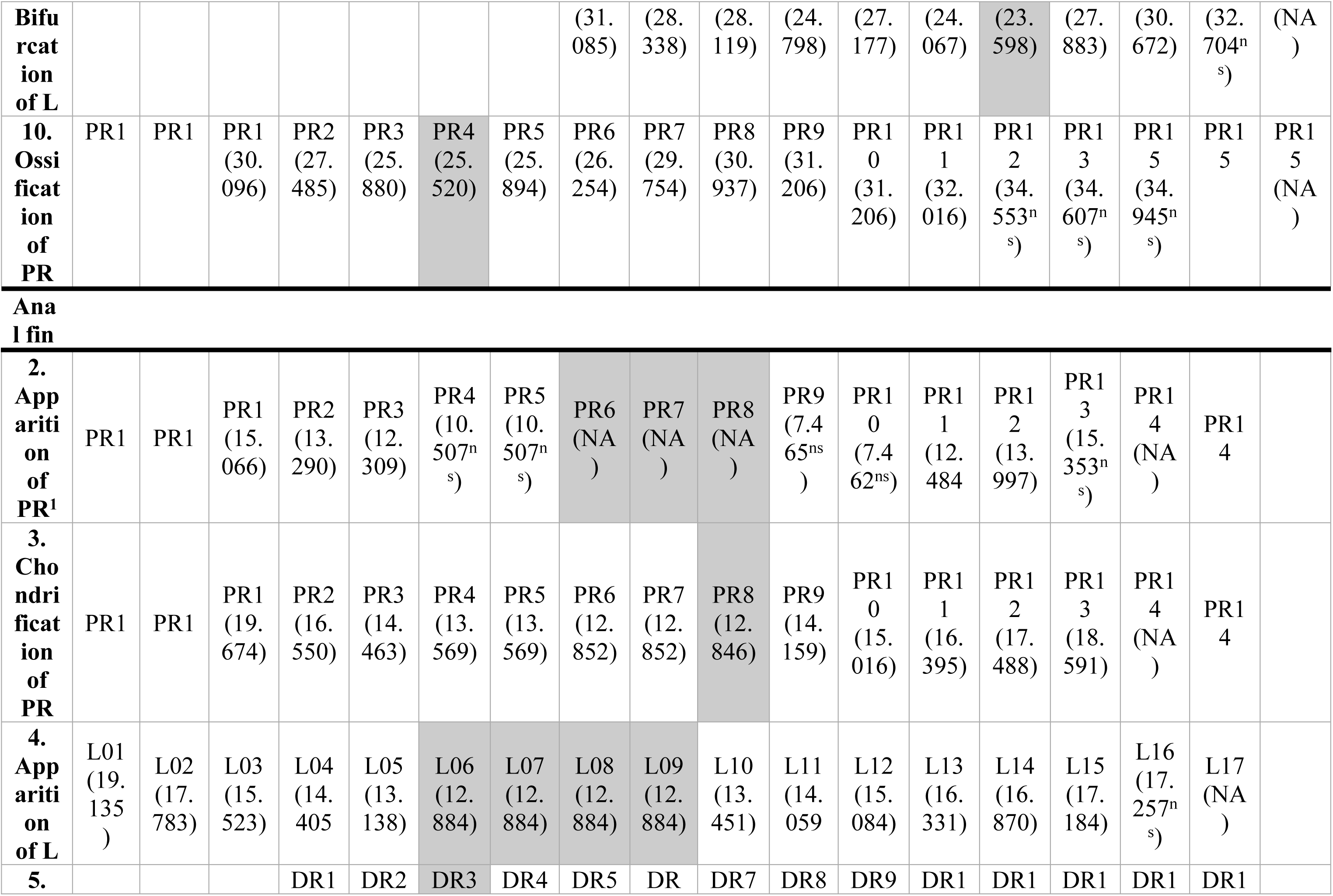
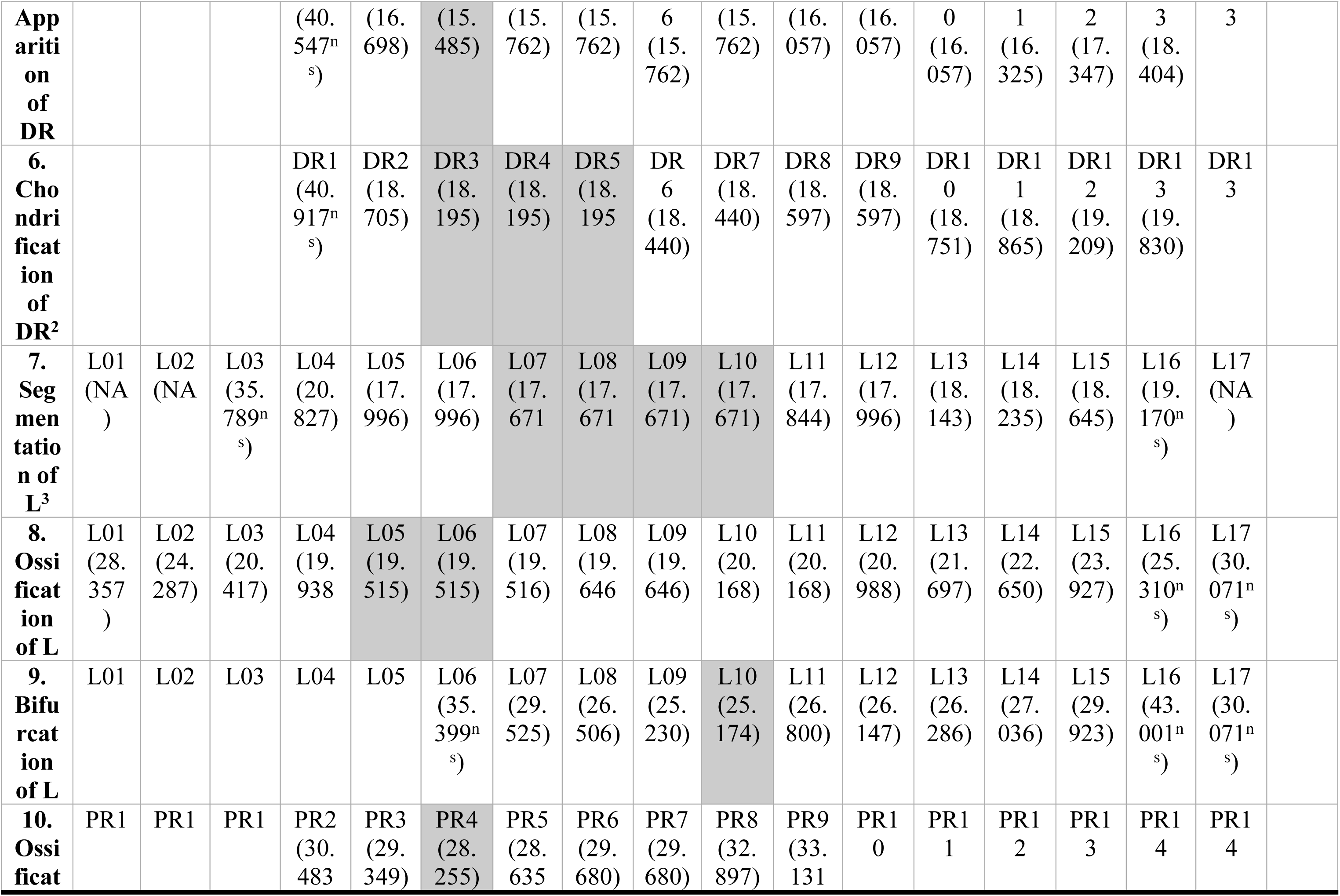
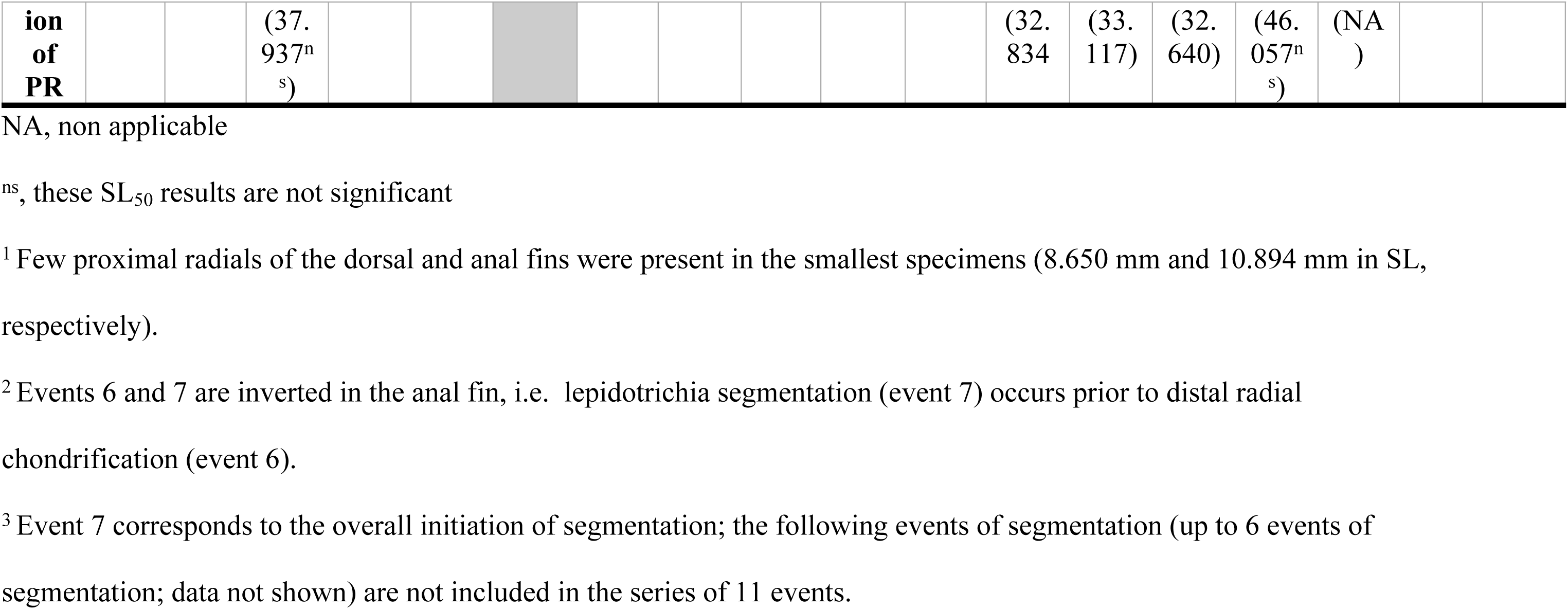
Values of the SL_50_ for the serial skeletal elements, i.e. the proximal radials (PR), the distal radials (DR) and the lepidotrichia (L) during nine out of eleven developmental events of the developmental sequence of the dorsal and anal fins of *Oncorhynchus*. See Nomenclature for the complete list of events. The serial skeletal elements are ordered from the anterior to the posterior of the fins and their column is representative of their position in the fin (see Fig 1). A PR or DR is found in more than one column when more than one L is articulated with it. The sites of initiation of the different events are identified with a grey shading.

The apparition of proximal radials (event 2) begins before hatching. Few proximal radials, located centrally, are present in the dorsal and anal fins of all the specimens examined, even in the smallest pre-hatching specimens (8-10-mm SL), thus, no SL_50_ were obtained for these elements (Table 2; Fig 3A). These proximal radials are interpreted as the initiation sites of the development of the dorsal (PR 5-9) and anal (PR 6-8) fin endoskeletons. These sites are congruent in the dorsal and anal fins (Table 2; Fig 3A). In both fins, proximal radials appear by proceeding bilaterally from the initiation site (Table 3). This indicates a bidirectional pattern for the development of the proximal radials; the most peripherally located proximal radials are the last to appear. The relative developmental sequences are not simultaneous but are significantly correlated between fins (Table 4). All following events occur after hatching.

**Fig 3.**
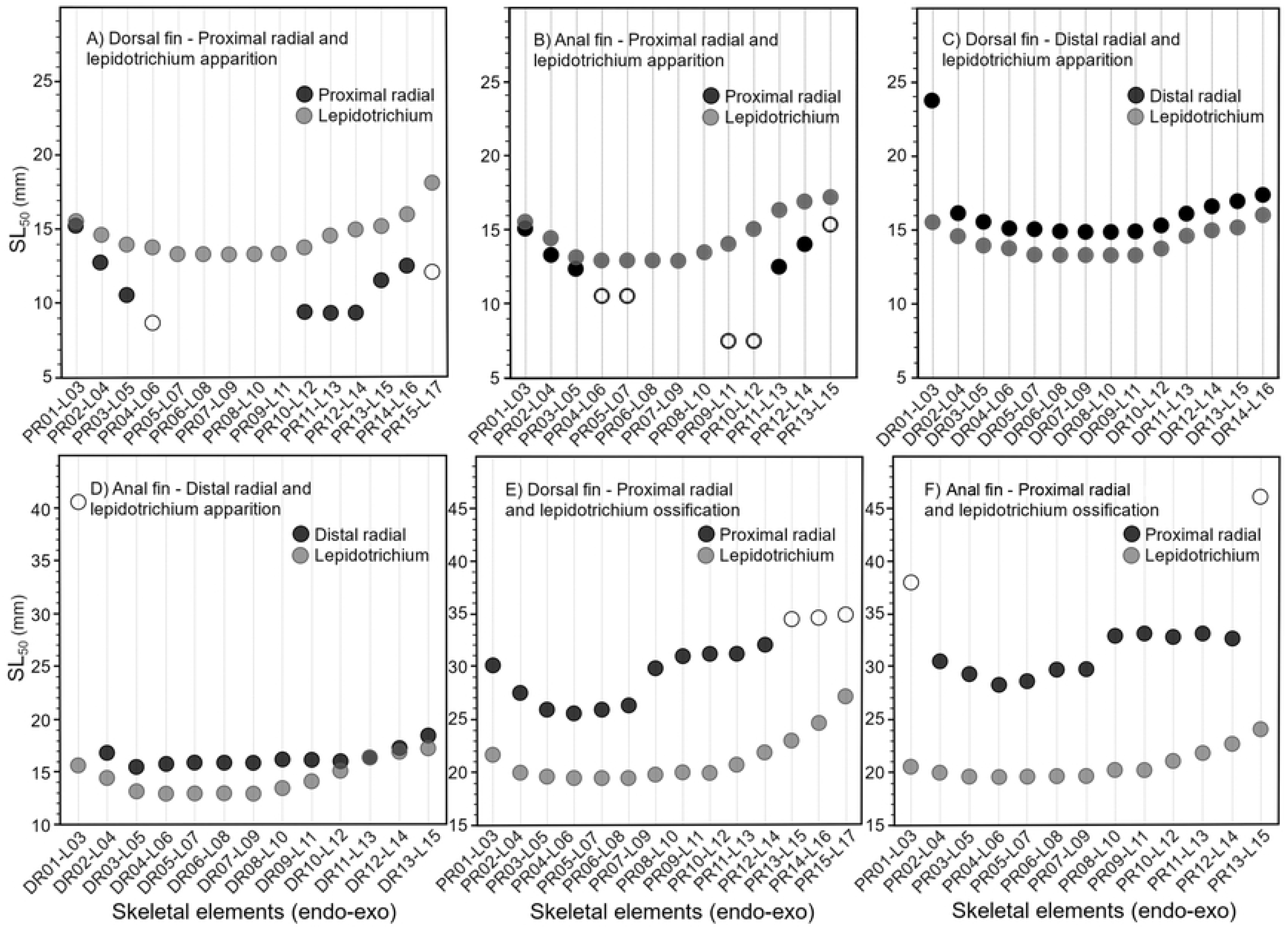
Comparisons between nine out of eleven developmental sequences for the dorsal (black) and anal (gray) fins of the rainbow trout (*Oncorhynchus mykiss*). See Nomenclature for the complete list of events. Skeletal elements are ordered from anterior to posterior. Filled and empty symbols represent significant and non-significant results for SL50, respectively.

**Table 3.** Directions of developmental sequences of ten out of eleven events for the dorsal (D) and anal (A) fins in living and extinct osteichthyans. See Nomenclature for the complete list of events.

| Events | *Oncorhynchus* |  | *Elonichthys* |  | *Miguashaia* |  | *Rhabdoderma* |  | *Quebecius* |  | *Dipterus* |  | *Eusthenopteron* |  |
| --- | --- | --- | --- | --- | --- | --- | --- | --- | --- | --- | --- | --- | --- | --- |
|  | D | A | D | A | D | A | D | A | D | A | D | A | D | A |
| 2. Apparition of proximal radials | B | B | N<br>A<sup>1</sup> | N<br>A<sup>1</sup> | N<br>A<sup>1</sup> | N<br>A<sup>1</sup> | NA<br>1 | NA<br>1 | N<br>A<sup>1</sup> | N<br>A<sup>1</sup> | N<br>A<sup>1</sup> | N<br>A<sup>1</sup> | NA<br>1 | NA<br>1 |
| 3. Chondrification of proximal radials | B | B | N<br>A<sup>1</sup> | N<br>A<sup>1</sup> | N<br>A<sup>1</sup> | N<br>A<sup>1</sup> | NA<br>1 | NA<br>1 | N<br>A<sup>1</sup> | N<br>A<sup>1</sup> | N<br>A<sup>1</sup> | N<br>A<sup>1</sup> | NA<br>1 | NA<br>1 |
| 4. Apparition of lepidotrichia | B | B | N<br>A<sup>1</sup> | N<br>A<sup>1</sup> | N<br>A<sup>1</sup> | N<br>A<sup>1</sup> | NA<br>1 | NA<br>1 | N<br>A<sup>1</sup> | N<br>A<sup>1</sup> | N<br>A<sup>1</sup> | N<br>A<sup>1</sup> | NA<br>1 | NA<br>1 |
| 5. Apparition of distal radials | B | B | N<br>A<sup>1</sup> | N<br>A<sup>1</sup> | N<br>A<sup>1</sup> | N<br>A<sup>1</sup> | NA<br>1 | NA<br>1 | N<br>A<sup>1</sup> | N<br>A<sup>1</sup> | N<br>A<sup>1</sup> | N<br>A<sup>1</sup> | NA<br>1 | NA<br>1 |
| 6. Chondrification of distal radials | B | B | N<br>A<sup>1</sup> | N<br>A<sup>1</sup> | N<br>A<sup>1</sup> | N<br>A<sup>1</sup> | NA<br>1 | NA<br>1 | N<br>A<sup>1</sup> | N<br>A<sup>1</sup> | N<br>A<sup>1</sup> | N<br>A<sup>1</sup> | NA<br>1 | NA<br>1 |
| 7. Segmentation of lepidotrichia | B | B | B | B | B | B | B | B | ? | ? | B | B | B | B |
| 8. Ossification of lepidotrichia | B | B | ? | ? | ? | ? | B | B | ? | ? | ? | ? | ? | ? |
| 9. Bifurcation of lepidotrichia | B | B | N<br>A<sup>2</sup> | N<br>A<sup>2</sup> | B | B | NA<br>2 | NA<br>2 | B | B | B | B | B | B |
| 10.<br>Ossificati<br>on of<br>proximal<br>radials | B | B | ? | ? | N<br>A<sup>3</sup> | N<br>A<sup>3</sup> | ? | ? | ? | ? | ? | ? | NA<br>3 | NA<br>3 |
| 11.<br>Ossificati<br>on of<br>distal<br>radials | NA<br>4 | NA<br>4 | B-A<br>P<sup>5</sup> | B-A<br>P<sup>5</sup> | N<br>A<sup>3</sup> | N<br>A<sup>3</sup> | ? | ? | ? | ? | ? | ? | PA | PA |

B, Bidirectional; AP, Antero-posterior; PA, Postero-anterior
?, the skeletal elements and/or sequence are unknown
NA: non-applicable
<sup>1</sup> This event is not observable in fossil taxa
<sup>2</sup> Bifurcation is unknown
<sup>3</sup> Only one endoskeletal element (for *Miguashaia*, the condition is unknown for the anal fin, however, the dorsal fin has at least three distal radial and one large basal plate)
<sup>4</sup> The ossification occurs after 100 dph.
<sup>5</sup> Data suggest an antero-posterior direction but a bidirectional direction is possible (see the text)

**Table 4.** Spearman correlations between developmental sequences of (a) the dorsal and anal fins and (b) fin exoskeleton and endoskeleton of the dorsal (D) and anal (A) fins of *O. mykiss* to validate the presence of the DAFPM and the EEM respectively. See Nomenclature for the complete list of events.

| Events | n | $r_s$ | $P$ |
| --- | --- | --- | --- |
| **Dorsal and anal fin patterning module** |  |  |  |
| 2. Proximal radial differentiation | 8 | 0.8553628 | <0.01 |
| 3. Proximal radial chondrification | 13 | 0.9185589 | <0.001 |
| 4. Lepidotrichia differentiation | 15 | 0.8604663 | <0.001 |
| 5. Distal radial differentiation | 12 | 0.6288943 | <0.05 |
| 6. Distal radial chondrification | 12 | 0.9019622 | <0.001 |
| 7. Lepidotrichia segmentation | 12 | 0.6691664 | <0.05 |
| 8. Lepidotrichia ossification | 15 | 0.9576292 | <0.001 |
| 9. Lepidotrichia bifurcation | 8 | -0.40476 | ns |
| 10. Proximal radial ossification | 10 | 0.9268293 | <0.001 |
| 11. Distal radial ossification | NA | NA | NA |
| **Endoskeleton and exoskeleton module** |  |  |  |
| Events 2 and 4: Proximal radial and lepidotrichia differentiation (D) | 14 | 0.903 | < 0.001 |
| Events 2 and 4: Proximal radial and lepidotrichia differentiation (A) | 12 | 0.730 | < 0.01 |
| Events 4 and 5 Lepidotrichia and distal radial differentiation (D) | 14 | 0.972 | < 0.001 |
| Events 4 and 5 Lepidotrichia and distal radial differentiation (A) | 12 | 0.891 | < 0.001 |
| Events 8 and 10 Lepidotrichia and proximal radial ossification (D) | 11 | 0.854 | < 0.001 |
| Events 8 and 10 Lepidotrichia and proximal radial ossification (A) | 11 | 0.839 | < 0.01 |

NA, non applicable: The ossification of distal radials occurs after 100 dph
ns, non significant

The chondrification of the proximal radials (event 3) quickly follows their appearance. In both fins, the chondrification sequences show similar patterns as to their sequences of apparition (S1 Table); chondrification starts from the same initiation site (i.e., the most centrally located proximal radials) and further proceeds bidirectionally (Table 3). The bidirectional sequences of chondrification are significantly correlated and are almost simultaneous between dorsal (PR 4-9) and anal (PR 8) fins (Fig 3B, Table 4).

Lepidotrichia appear (event 4) slightly after the differentiation of the first proximal radial and subsequently articulate with the distal radials. The initiation site of the lepidotrichia is similar in both fins (L7-11 in the dorsal fin and L6-9 in the anal fin) and corresponds in position to the early apparition of the proximal radials (S1 Table). Similar to proximal radials, new lepidotrichia appear by proceeding bidirectionally from the initiation site (Table 2). The sequences of apparition of lepidotrichia are highly correlated between fins and almost simultaneous (Fig 3C, Table 4).

The first distal radials (event 5) to appear differ in position between the dorsal (DR6-9) and anal (DR3) fins (Table 2). In both fins, the sequences of apparition proceed bidirectionally from the initiation site. The sequences are significantly correlated (Fig 3D, Table 4).

The chondrification of distal radials (event 6) follows rapidly, starting from similar initiation sites, DR5 in the dorsal fin and DR3-5 in the anal fin (Table 2). Starting from the initiation site, the chondrification sequences proceed bidirectionally and are almost simultaneous as well as highly correlated between fins (Fig 3E, Table 4).

Lepidotrichia grow by distal addition of new segments, a process called segmentation (event 7). Through growth, five and six segmentation events occur in the anal and dorsal fins, respectively. The lepidotrichia with the higher numbers of segments in most specimens are the L10-14 in the dorsal fin and L8-10 in the anal fin and the number of segments on the adjacent lepidotrichia decreases in a bidirectional pattern. The first segmentation event occurs slightly posteriorly in the dorsal fin (L12) comparatively to the anal fin (L7-10) (Fig 3F). The site for the initiation of segmentation corresponds to the lepidotrichia with the highest number of segments, suggesting that the longest lepidotrichia, in terms of number of segments, are the first lepidotrichia to segment in the developmental sequence (Fig 1). All sequences of segmentation are bidirectional (Table 3). The sequences of the first segmentation are almost simultaneous and significantly correlated between fins (Fig 3F, Table 4).

The initiation site for the ossification of lepidotrichia (event 8) is similar in both fins (L6-8 in the dorsal fin and L5-6 in the anal fin) (Fig 3G). Sequences of ossification are bidirectional, simultaneous, and highly correlated (Tables 3 and 4). There are small, positional differences for the initiation site of apparition, segmentation and ossification of the lepidotrichia (Fig 3G). Nevertheless, sequences of apparition are significantly correlated with sequences of segmentation of the lepidotrichia in the dorsal (0.670; p<0.01) and anal (0.768; p<0.01) fins, while the results differ between sequences of segmentation and ossification of the lepidotrichia (dorsal, 0.098; p>0.05; anal, 0.485; p>0.05) (Table 4).

The bifurcation of the lepidotrichia (event 9) is initiated at different positions in the dorsal (L14) and anal (L10) fins (Fig 3H). The bidirectional sequences of bifurcation between fins are not significantly correlated and not simultaneous (Fig 3H, Table 4). Generally, in the dorsal fin, the lepidotrichia located at the initiation site show more proximal bifurcations comparatively to the lepidotrichia located bilaterally of the initiation site; this pattern is not as clear in the anal fin. A single order of bifurcation is present.

The ossification of proximal radials (event 10) is initiated from PR4, located in the same anterior portion of both fins (Fig 3I). In both fins, the sequences of ossification proceed in a bidirectional direction, and are highly correlated, but not simultaneous (Fig 3I, Table 4).

All nine developmental events analysed show a certain degree of congruence between the dorsal and anal fins (e.g., similar initiation site and/or similar direction) and eight events are significantly correlated between fins (events 2-8 and 10; Table 4).

Moreover, in congruence with the one-to-one relationship observed between radials and lepidotrichia in both fins, the first lepidotrichia to appear are related to the first apparition of proximal and distal radials (Fig 4); this is corroborated by the strong correlations between sequences of apparition of the endoskeletal and exoskeletal elements within each fin (Table 4). Highly significant correlations are also found for sequences of ossification of the endoskeleton and exoskeleton within both fins (Fig 4, Table 4).

**Fig 4.**
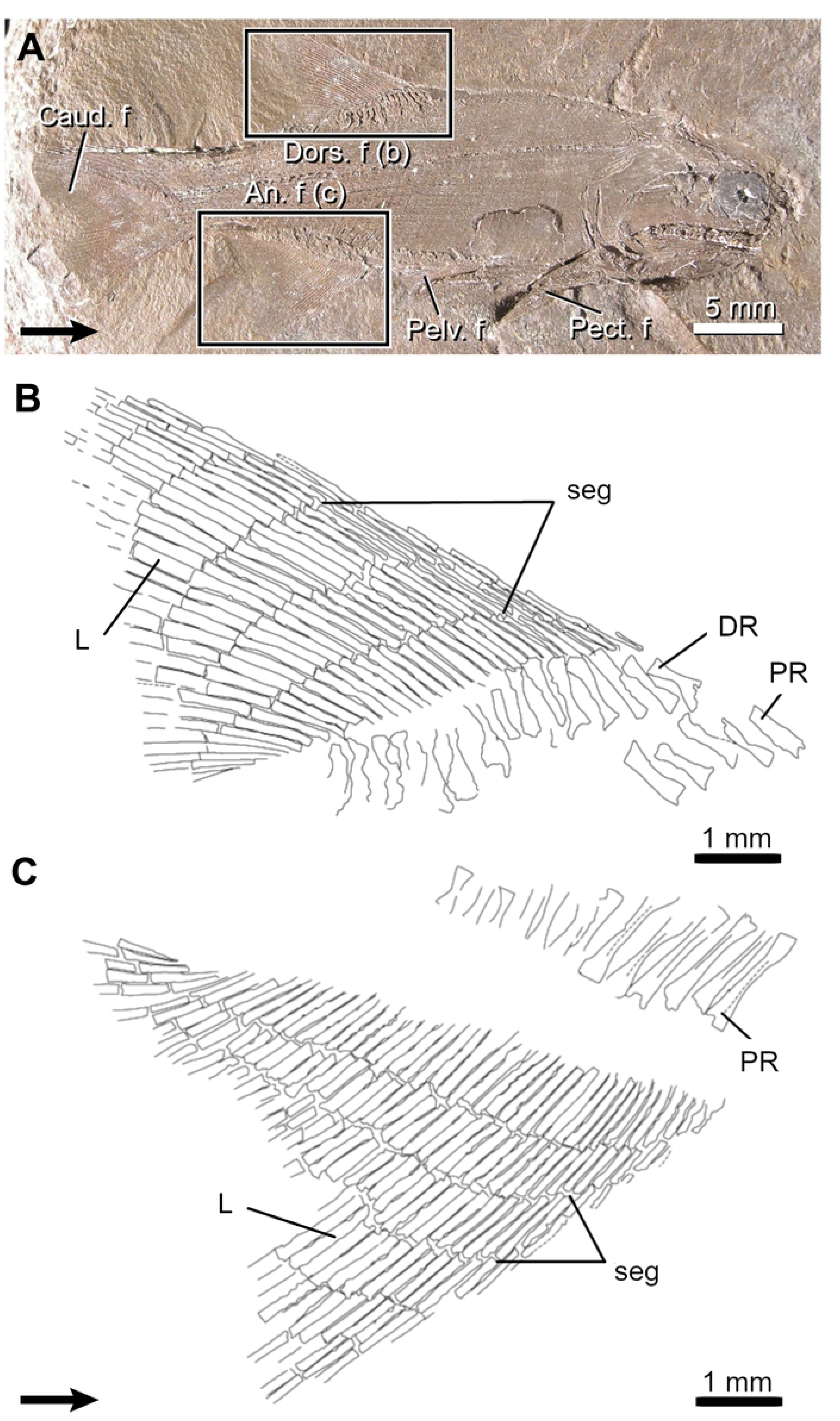
Comparisons between developmental sequences (apparition and ossification) of the endoskeleton (radials; black) and exoskeleton (lepidotrichia; grey) of the dorsal (A, C, E) and anal fins (B, D, F) of the rainbow trout (*Oncorhynchus mykiss*). Skeletal elements are ordered from anterior to posterior. Filled and empty symbols represent significant and non-significant results for SL50 respectively.

### Developmental patterning in the dorsal and anal fins of fossil osteichthyans

For the six fossil osteichthyans surveyed (Table 1, S1 Table), only the events dealing with the lepidotrichia [i.e., segmentation (event 7), ossification (event 8), and bifurcation (event 9)] and the ossification of endochondral elements [i.e., proximal radials (event 10) and distal radials (event 11)] are available to study due to the nature of fossils and the rarity of fossilized ontogenies. Meristic counts (Table 1) are based on mean values from all specimens examined for each taxon, including both juveniles and adults whenever possible. Congruence of events between the dorsal and anal fins, and between the endoskeleton and exoskeleton are inferred based on initiation sites, and when available, the direction of sequences.

#### Actinopterygians

##### Elonichthys peltigerus

The dorsal and anal fins of *Elonichthys* are of similar shape but differ slightly in size (Table 1; Fig 5) [47]. The dorsal fin is composed of 16-19 distal radials whereas the anal fin has 19-23 proximal radials. The general relationship between radials and lepidotrichia is 1:2, with some supernumerary lepidotrichia occurring in the anterior and posterior margins.

**Fig 5.**
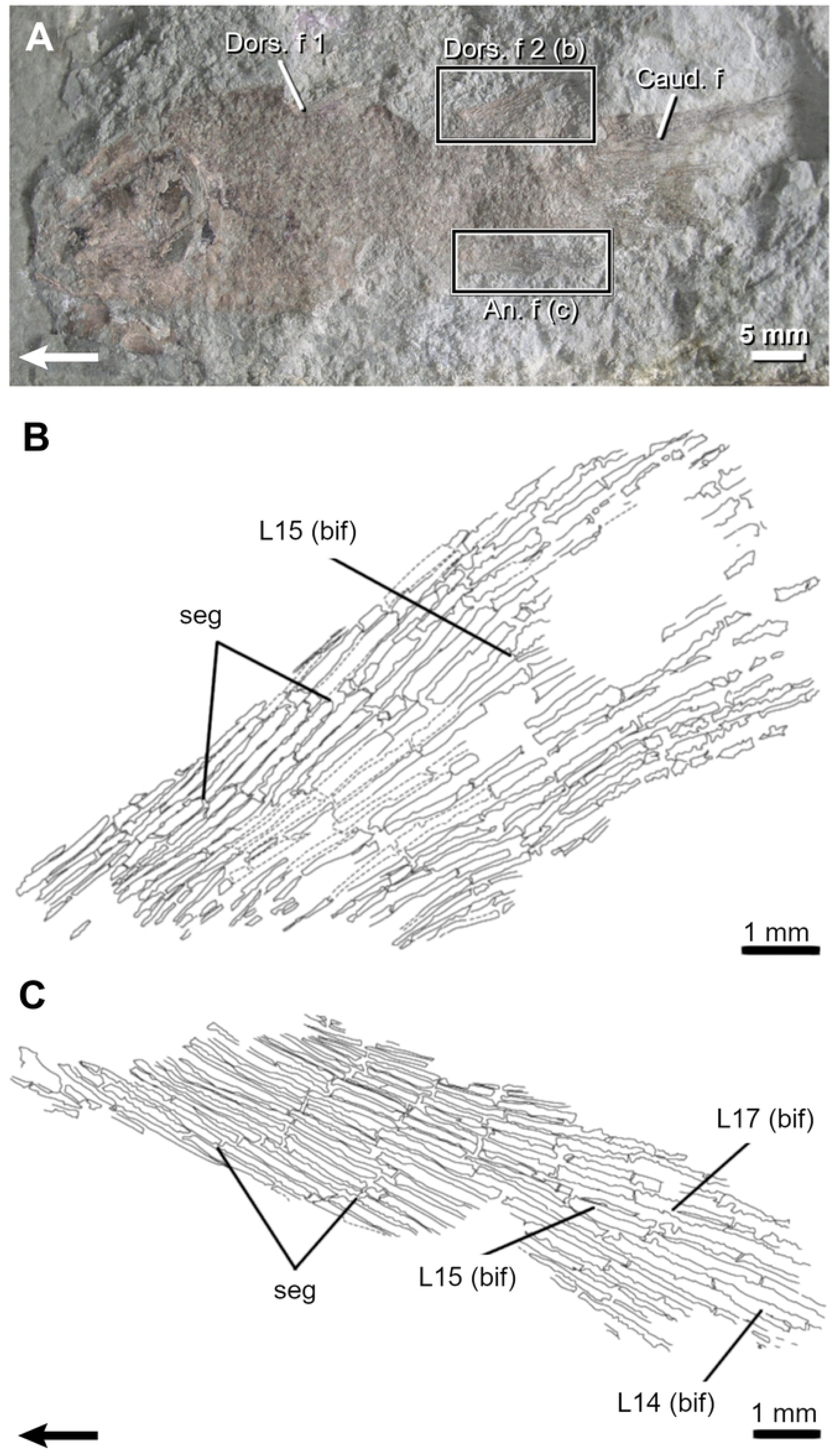
The Carboniferous actinopterygian *Elonichthys peltigerus*. A, Fossil specimen (FMNH PF 7502) and drawings of its dorsal (B) and anal (C) fins. Arrows point anteriorly. An. f, anal fin; caud. f, caudal fin; Dors. f, dorsal fin; DR, distal radial; L, lepidotrichia; Pect. f, pectoral fin; Pelv. f, pelvic fin; PR, proximal radial; seg, segmentation.

Lepidotrichia are the first structures to ossify in both fins. Segmented lepidotrichia (event 7) are present in all specimens. The number of segments varies little with respect to SL (from 4-5 to 6-7 segments per lepidotrichium). In both fins, lepidotrichia from the anterior part of the fin are usually the longest (L7-15 in the dorsal and L10-13 in the anal fin) and articulate with distal radials 5-6, which seem among the first ones to ossify (see below). The number of segments decreases bilaterally from these lepidotrichia.

Lepidotrichia are already numerous in the smallest specimen (38 dorsal and 46 anal lepidotrichia); therefore, it is not possible to infer a sequence of ossification (event 8). None of the specimens show bifurcated lepidotrichia (event 9).

Ossification sequences are difficult to reconstruct for the proximal (event 10) and distal radials (event 11). Proximal radials are visible at the anterior portion of the dorsal and anal fins; remaining radials are probably hidden under the scale cover. Distal radials have only been clearly identified in the dorsal fin (FMNH PF 7502) (Fig 5). When present, both ossified proximal and distal radials do not reach the posterior margin of both fins.

#### Actinistians

##### Miguashaia bureaui

The second dorsal and anal fins of *Miguashaia* are overly similar in size and shape (Table 1; Figs 6 and 7) with narrow, rectangular bases and pointed anterior corners ([22], fig. 1B). Only the lepidotrichia can be described (27-28 in the dorsal fin and ca. 25 in the anal fin), a single specimen (MHNM 06-1809) partially shows the radials articulating with the second dorsal basal plate.

**Fig 6.**
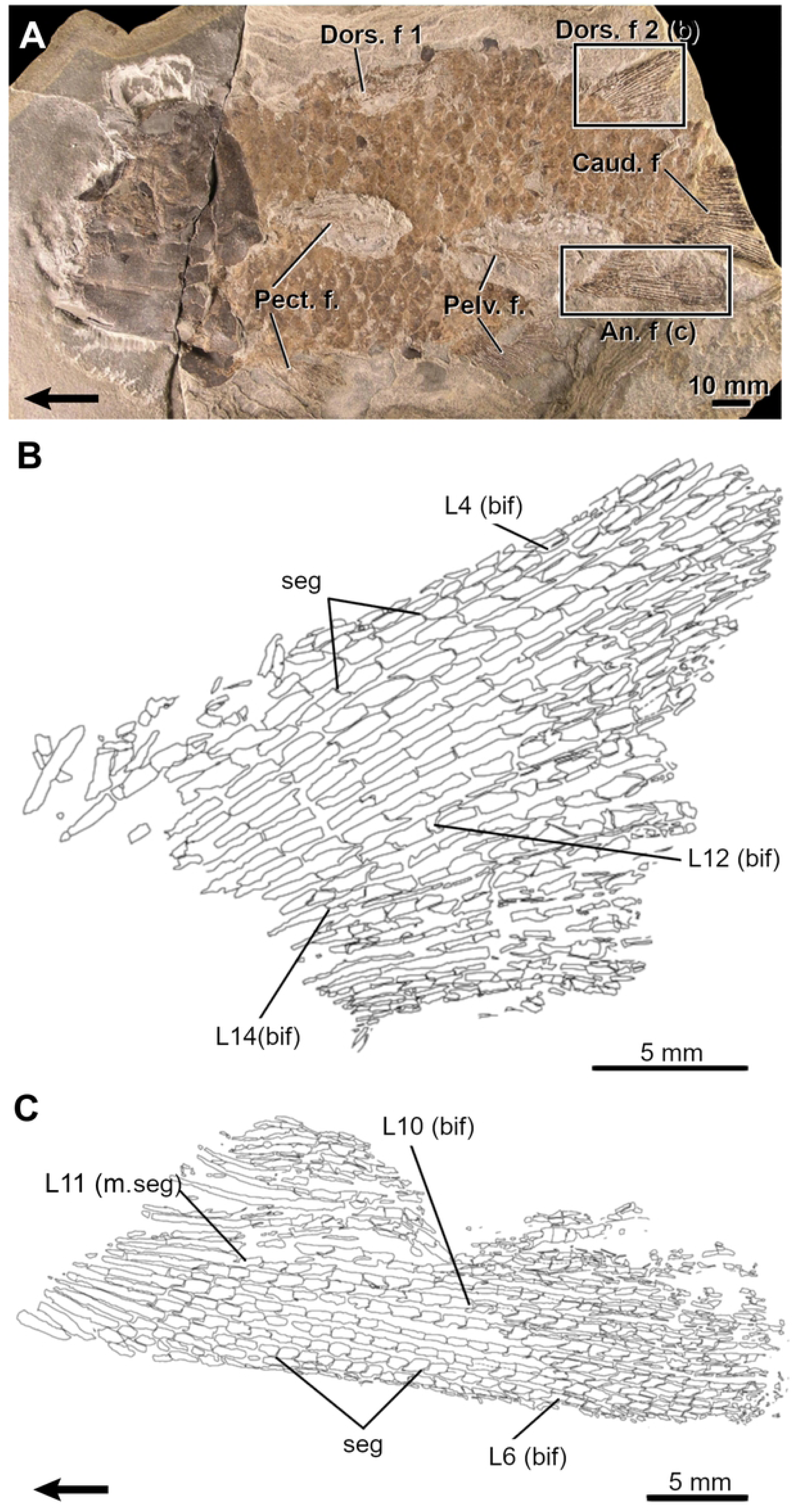
The Devonian coelacanth *Miguashaia bureaui*. A, Juvenile specimen (ULQ 120) and drawings of its second dorsal (B) and anal (C) fins. Arrows point anteriorly. An. f, anal fin; bif, bifurcation; Caud. f, caudal fin; Dors. f, dorsal fin; L, lepidotrichia; seg, segmentation.

**Fig 7.**
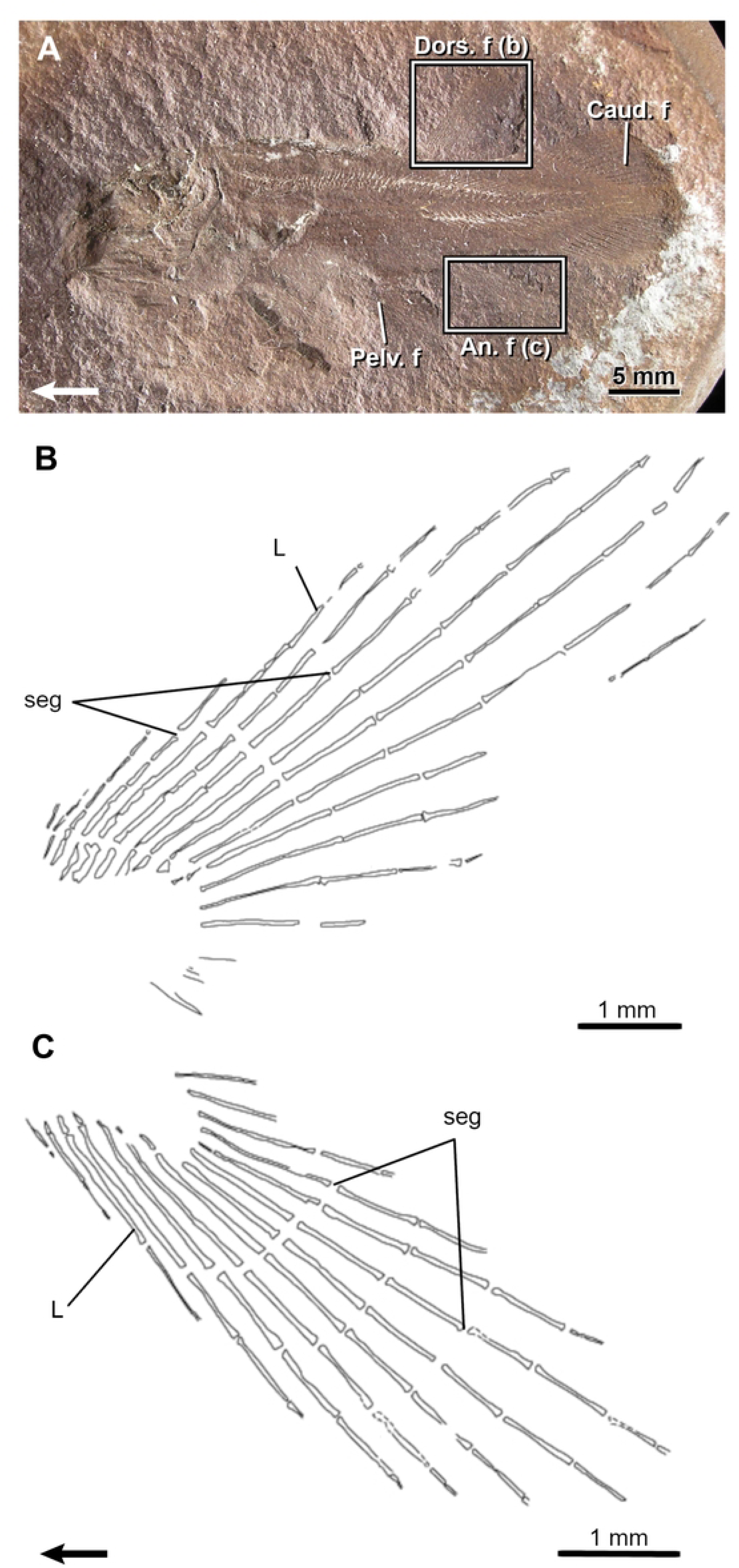
The Devonian coelacanth *Miguashaia bureaui.* A, Adult specimen (MHNM 06-41) and drawings of its second dorsal (B) and anal (C) fins. Note the occurrence of merging segments (m.seg) at the base of the fins. Arrows point anteriorly. An. f, anal fin; bif, bifurcation; Caud. f, caudal fin; Dors. f, dorsal fin; L, lepidotrichia; Pect. f, pectoral fin; Pelv. f, pelvic fin; PR, proximal radial; seg, segmentation.

Numerous segmentations (up to five) are visible in the lepidotrichia (event 7) of the dorsal and anal fins of the smallest specimen (MHNM 06-1633, 64.5 mm TL). The longest lepidotrichia are L4-7 in the dorsal fin and L5-8 in the anal fin in specimen MHNM 06-41 (Fig 7) and the number of segments decreases gradually bilaterally from these sites. The basal proximal segment is always longer than the others in all specimens examined. In specimen MHNM 06-41, the segment immediately distal to the first proximal segment appears to be half-fused with this basal segment (Fig 7; L11, m.seg), thus evidencing that the increase in length of the basal segment during growth is the result of its merging with other proximal segments.

Ossified lepidotrichia are already numerous (28 dorsal and ca. 22 anal lepidotrichia) in the smallest specimen (MHNM 06-1633); it is thus not possible to infer an ossification sequence (event 8).

Few bifurcations (event 9) are seen on L15 and L14-17, respectively in the dorsal and anal fins of the small specimen ULQ 120 (85 mm TL; Fig 6). The most proximal bifurcations are positioned approximately between L12-15 in all specimens (Figs 6 and 7). Bifurcations are present anteriorly and posteriorly to L12-15 in large specimens.

Bifurcations are gradually displaced distally anterior to the most proximally bifurcated lepidotrichium (L15) resulting in a higher proportion of proximally bifurcated lepidotrichia in the posterior portion of the fins. Up to three orders of bifurcation have been observed in specimen MHNM 06-494 (400 mm TL).

Ossified endoskeletal supports (events 10 and 11) are poorly documented. A single basal plate in the first dorsal fin is present in one partial specimen (MHNM 06-1232), while the basal plate of the second dorsal fin shows articulating surfaces for three radials, the posterior one being well-preserved (MHNM 06-1809).

##### Rhabdoderma exiguum

The second dorsal and anal fins of *Rhabdoderma* are almost identical in size and shape (Table 1, Fig 8). They display a narrow-based fan-like outline and, as for *Miguashaia*, only the lepidotrichia are preserved.

**Fig 8.**
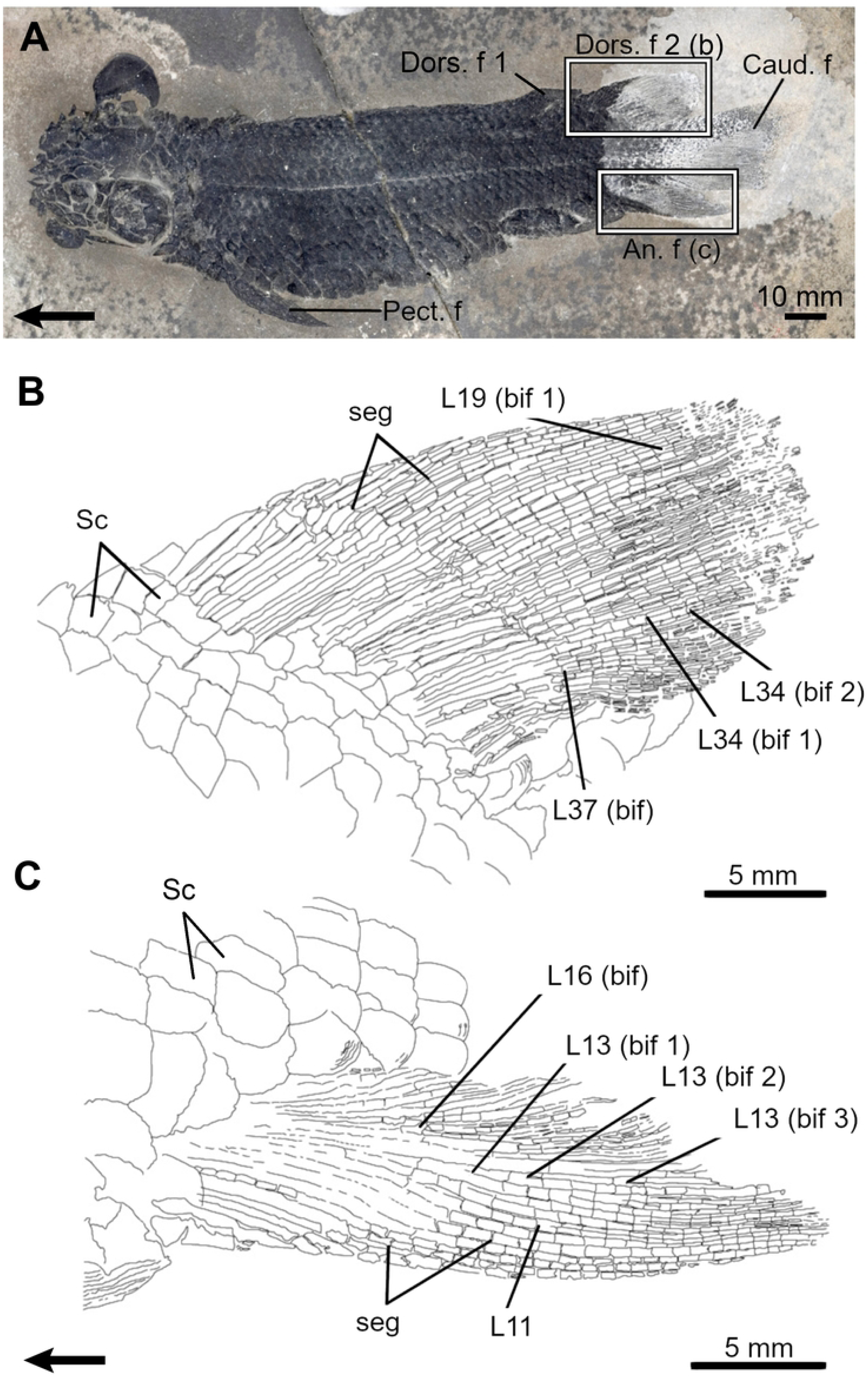
The Carboniferous coelacanth *Rhabdoderma exiguum.* A, Small specimen (FMNH PF 9954) and drawings of its second dorsal (B) and anal (C) fins. Arrows point anteriorly. An. f, anal fin; Caud. f, caudal fin; Dors. f, dorsal fin; L, lepidotrichia; Pelv. f, pelvic fin; seg, segmentation.

All specimens show segmented lepidotrichia (event 7). The number of segments increases with SL (from 5-6 to 13-14 segments). In both fins, the longest lepidotrichia occur between L9-12 and the number of segments gradually decreases bilaterally from this site.

The number of lepidotrichia increases up to 21 in both fins during growth (event 8). Lepidotrichia at the anterior and posterior extremities are absent in the dorsal and anal fins of small specimens, suggesting that central lepidotrichia might have ossified before the anterior and posterior ones (Fig 8). This fin comparison between small (less than 21 lepidotrichia; e.g., FMNH PF7528; 33 mm SL) and larger specimens (21 lepidotrichia; e.g., FMNH PF7338; 48 mm SL) was done by lining up their respective longest lepidotrichia.

None of the specimens show bifurcation of the lepidotrichia (event 9), and no ossified endoskeletal supports are preserved in embryos and larvae examined (events 10 and 11).

#### Porolepiforms

##### Quebecius quebecensis

The second dorsal and anal fins of *Quebecius* have a similar size and shape (Table 1) [48 (figs 1, 7), 49 (fig. 2)]. Endoskeletal supports remain unknown in this taxon and only the lepidotrichia can be described (between 30 and 40 lepidotrichia in both fins).

Segmented lepidotrichia (event 7) are already visible in the smallest specimen (MHNM 06-1474a); however, the number of segments is unclear in all specimens owing to preservation issues. Lepidotrichia are well developed, and their number is similar in the small as well as in the larger specimens (35 dorsal lepidotrichia and 34 anal lepidotrichia), it is thus not possible to infer an ossification sequence (event 8).

The first bifurcation (event 9) is seen in L22 of the anal fin (MHNM 06-1474a). The most proximal bifurcations occur between L18-21 in longer specimens, with other branched lepidotrichia located anteriorly and posteriorly to this area. In both fins, the posterior lepidotrichia display more proximal bifurcations than the anterior lepidotrichia.

#### Dipnoans

##### Dipterus valenciennesi

The second dorsal and anal fins of *Dipterus* differ in size and shape (Table 1; Fig 9), with the second dorsal fin being longer and higher than the somewhat pointed, leaf-shaped anal fin. Lepidotrichia are more numerous than their supporting radials, which are known to be grossly similar between fins with minor differences in terms of the number and shape [46].

**Fig 9.**
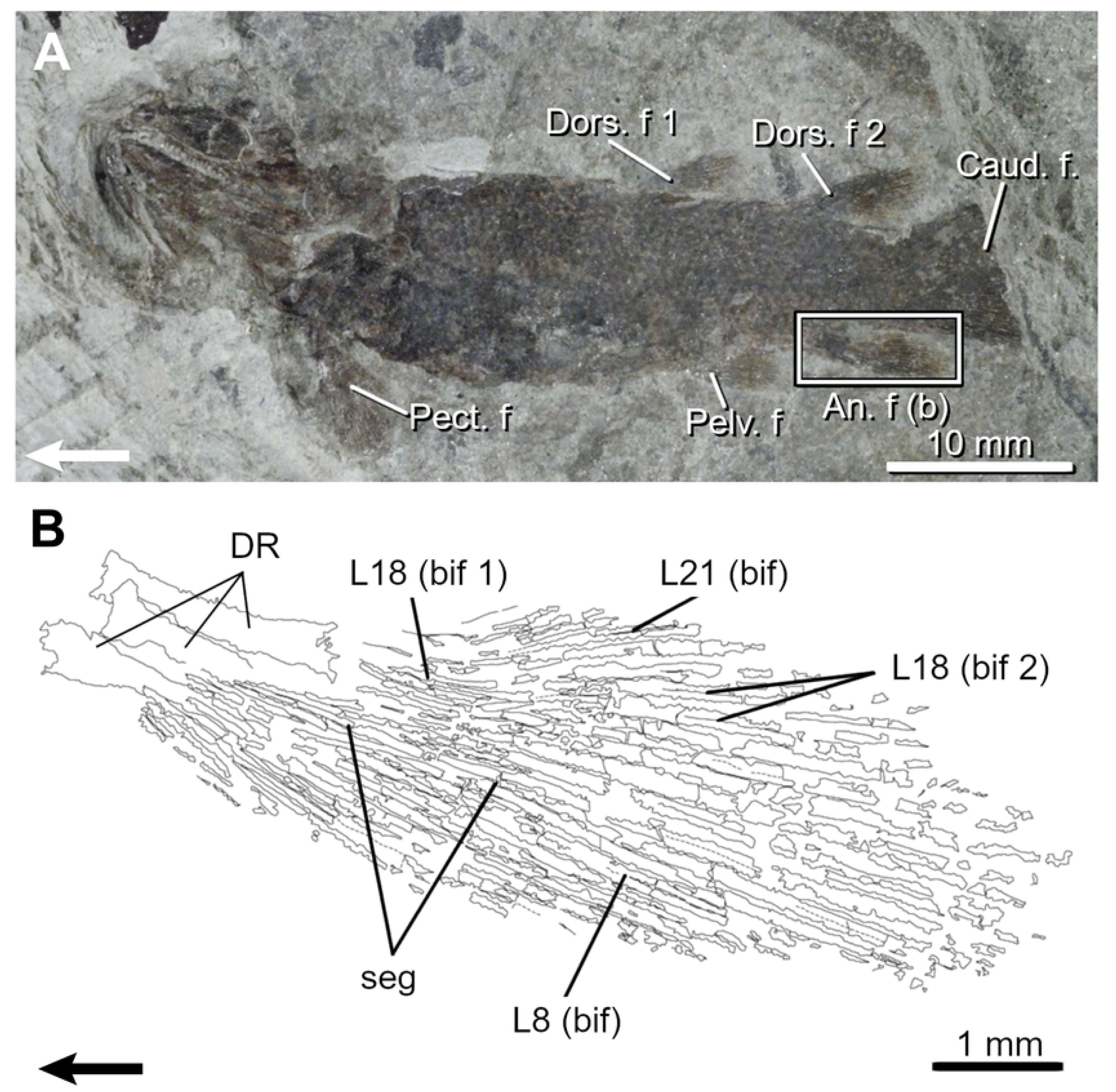
The Devonian dipnoan *Dipterus valenciennesi*. A, Fossil specimen of (BMNH P.22187) and drawings of its second dorsal (B) and anal (C) fins. Note the development of scales (Sc) covering the proximal portion of the lepidotrichia. Arrows point anteriorly. An. f, anal fin; bif, bifurcation; Caud. f, caudal fin; Dors. f, dorsal fin; L, lepidotrichia; Pect. f, pectoral fin; seg, segmentation.

Lepidotrichia show multiple segmentations (event 7). The longest lepidotrichia are not found at the same position in the dorsal (L16-26) and anal (L8-12) fins; these areas are interpreted as different initiation sites of segmentation. In both fins, the number of segments gradually decreases bilaterally from these sites. The basal segment is considerably long, comprising one third of each lepidotrichium total length.

Even the smallest specimens show numerous well ossified lepidotrichia (44 dorsal and 25 anal), it is thus not possible to infer a sequence of ossification (event 8).

Up to three orders of bifurcation (event 9) are present. Bifurcations are present between L14-18 and L39-42 in the dorsal fin and between L11-12 and L17-22 in the anal fin. Specimen BMNH P22187 (Fig 9A) is informative concerning the sequence of bifurcation: (1) two orders of bifurcation are present in the dorsal fin: 1^st^ order: L15-42 (bif 1), 2^nd^ order: L28-39 (bif 2) (Fig 9B), and three orders in the anal fin: 1^st^ order: L10-20 (bif 1); 2^nd^ order: L11-18 (bif 2); 3^rd^ order: L13-16 (bif 3) (Fig 9C); (2) the most proximal bifurcations are in L36-37 in the dorsal fin and in L13-16 in the anal fin (Figs 9B and 9C), and (3) bifurcated lepidotrichia are found anteriorly and posteriorly to these sites. Thus, in both fins, the posterior-most lepidotrichia show more proximal bifurcations whereas the anterior-most lepidotrichia are more distally branched.

Ossified endoskeletal supports (events 10 and 11) were not observed due to the scale cover (Sc, Figs 9B and 9C). Their sequence of ossification is unknown.

#### ‘Osteolepiforms’

##### Eusthenopteron foordi

The second dorsal and anal fins of *Eusthenopteron* are similar in size and shape (Table 1; Figs 10 and 11), displaying a narrow-based and posteriorly pointed profile [50]. The median fins display a broad basal plate on which generally three distal radials articulate, carrying numerous lepidotrichia (up to 25 in each fin).

**Fig 10.**
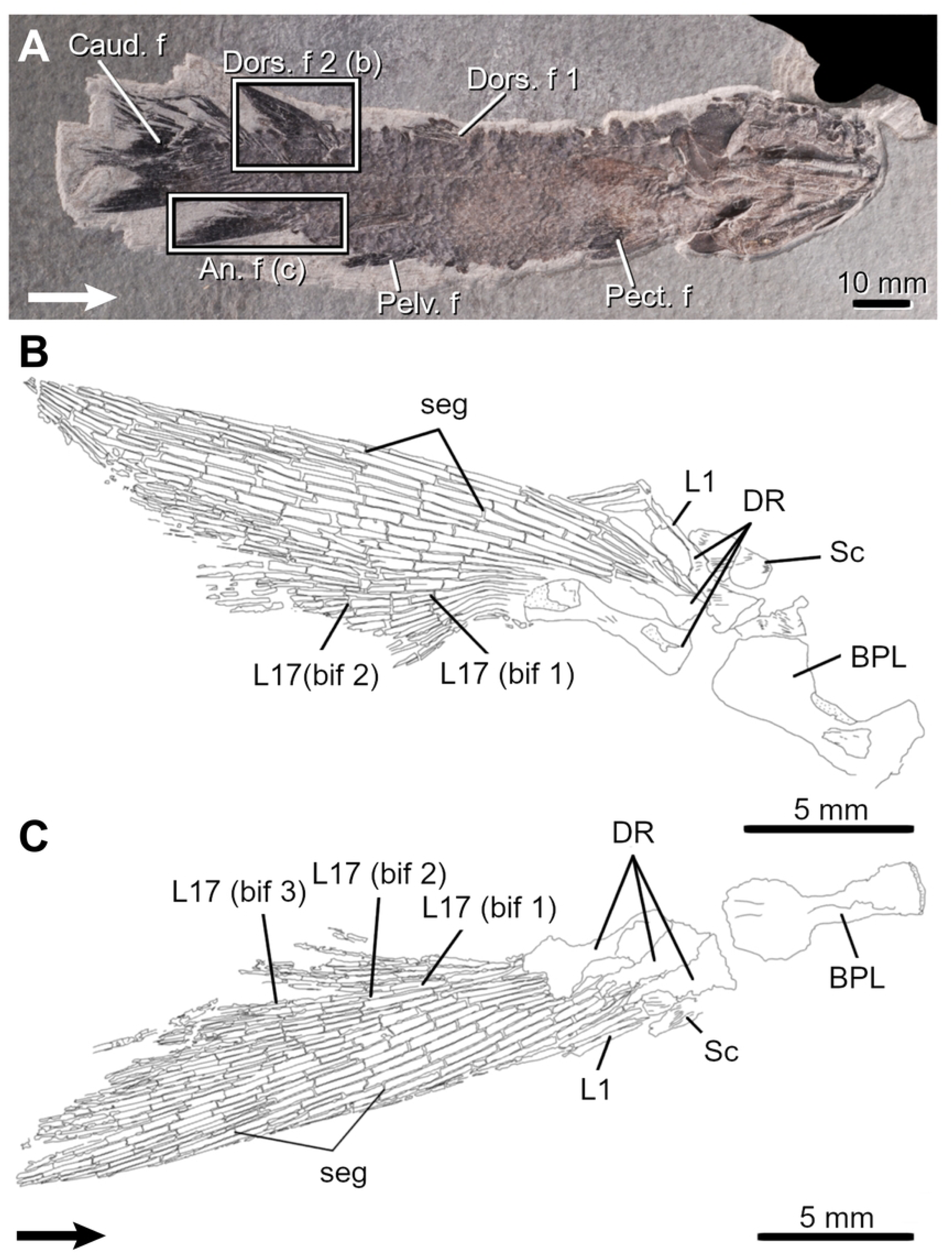
The Devonian ‘osteolepiform’ *Eusthenopteron foordi.* A, Very small specimen (MHNM 06-1754) and a drawing of its anal fin (B). Arrows point anteriorly. An. f, anal fin; bif, bifurcation; Caud. f, caudal fin; Dors. f, dorsal fin; DR, distal radial; L, lepidotrichia; Pect. f, pectoral fin; Pelv. f, pelvic fin; seg, segmentation.

**Fig 11.**
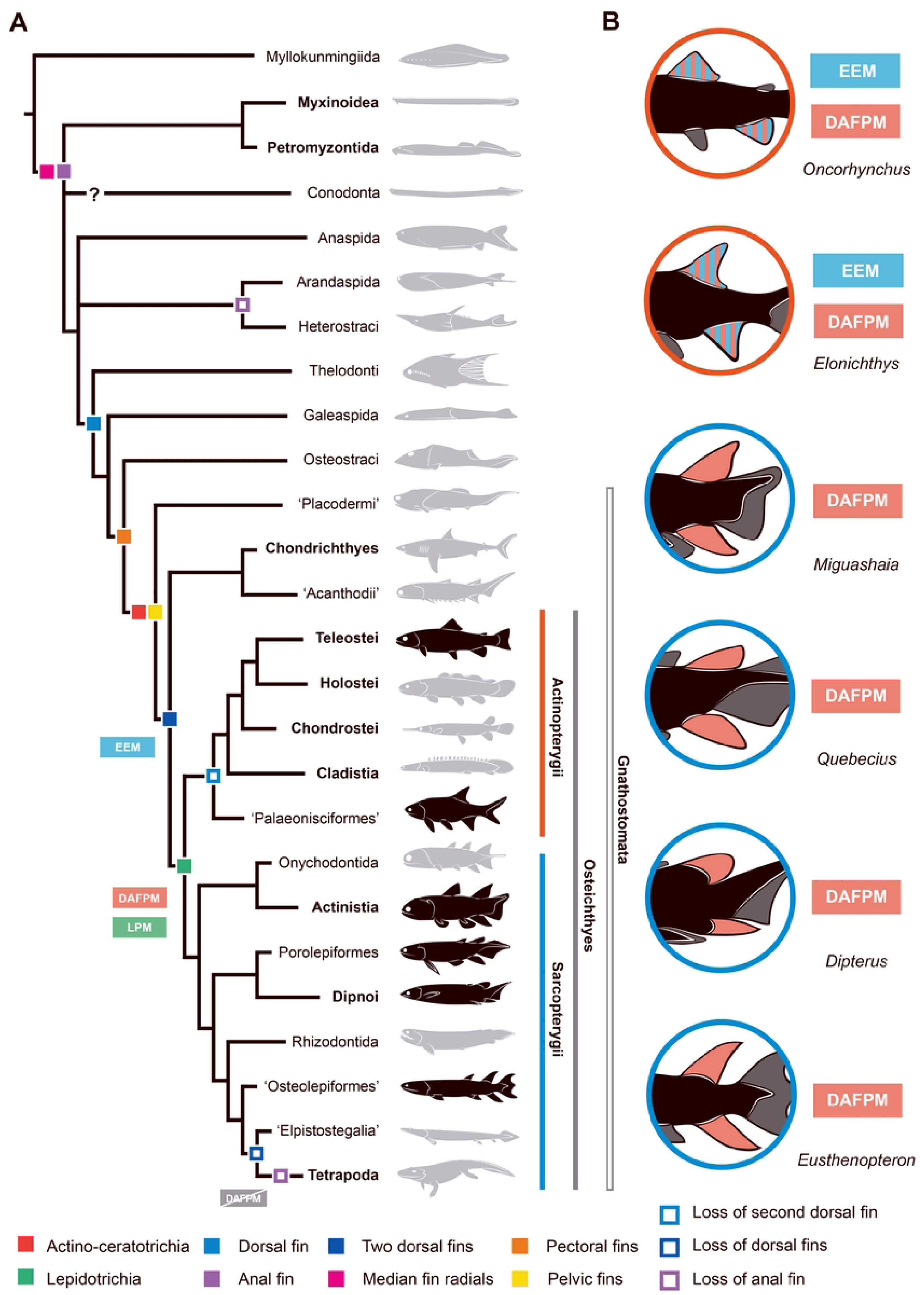
The Devonian ‘osteolepiform’ *Eusthenopteron foordi.* A, Small specimen (MHNM 06-1769) and drawings of its second dorsal (B) and anal (C) fins. Arrows point anteriorly. Note the preservation of a basal plate (BPL) and a reduced scale covering (Sc) at the base of the fins. An. f, anal fin; bif, bifurcation; Caud. f, caudal fin; Dors. f, dorsal fin; DR, distal radial; L, lepidotrichia; Pect. f, pectoral fin; Pelv. f, pelvic fin; PR, proximal radial; seg, segmentation.

The first segmentations (event 7) are seen in L11-14 in the dorsal fin and in L8-13 in the anal fin (MHNM 04-1293p10-Ef1; 40.8 mm SL). The position of these first segmented lepidotrichia is congruent with the location of the longest lepidotrichia in larger specimens. In these specimens, most lepidotrichia are segmented and the number of segments gradually decreases bilaterally from L8-14.

Lepidotrichia are the first structures to ossify in both fins (event 8); they are numerous in the smallest specimen (MHNM 04-1293p9-Ef1, 27.4 mm SL; 18 and 21 lepidotrichia in dorsal and anal fins, respectively) and the highest number of lepidotrichia is reached early (MHNM 06-1754; 49.3 mm SL; Fig 10), thus it is not possible to infer an ossification sequence.

Bifurcations (event 9) are restricted between L6-7 to L25 in large specimens and up to five orders of bifurcation are present in both fins. First bifurcations are present in L13 and 17 in the dorsal fin and in L18-19 in the anal fin (MHNM 06-1754; Fig 10B) Most proximal bifurcations and subsequent orders of bifurcation are initiated in this area (e.g., MHNM 06-1769: 2^nd^ order: L9-17 in the dorsal fin and L13-18 in the anal fin; 3^rd^ order: L15-17 in the anal fin; Figs 11B and 11C).

Radial ossification (DR; event 11) occurs prior to the ossification of the basal plate (event 10) (BPL, Figs 11B and 11C). A postero-anterior pattern of ossification for dorsal and anal distal radials is suggested because few small specimens (43-56-mm SL) show only the posterior, or the posterior and central radials in the dorsal or anal fins. The basal plate is first seen at 88.9-mm SL and 69.6-mm SL in the dorsal and anal fins, respectively.

## Discussion

Our study described a series of 11 skeletogenic events in the formation of the endo- and exoskeletal components of the dorsal and anal fins of the living actinopterygian *Oncorhynchus mykiss* between 5 days pre-hatching up to 100 days post-hatching. We also documented some of these events in one Carboniferous actinopterygian species, and five Devonian-Carboniferous sarcopterygian species. For the first time, we quantified the segmentation and bifurcation of lepidotrichia supporting a lepidotrichial patterning module.

The similarity and synchronicity of these developmental events between the dorsal and anal fins support the hypothesis that these two fins form a developmental module.

### Median fins in vertebrates

The fins of vertebrates can be described as membranous lateral outgrowths of the body walls reinforced internally by elongated elements, which can be of endoskeletal (e.g., radial bones) or dermal (e.g., fin rays) origin.

Median fins are present in stem vertebrates, such as myllokunmingiids, in the form of dorsal and ventral skin folds reminiscent of the median larval fin fold observed during the early ontogeny of more advanced fishes (e.g., [6,9,11,51]). The earliest ‘agnathans’ display well-developed median fins, which include, in most cases, a caudal fin and elongate dorsal and ventral fins. A separate anal fin has been confidently identified in petromyzontiforms, anaspids [52, 53], the anaspid-like *Euphanerops,* and in certain thelodonts (e.g., *Loganellia*) [54]. In *Euphanerops*, the anal fins are paired fins [55, 56].

The presence of an anal fin in hagfishes has been suggested in *Myxinikela* [57], but it is currently difficult to confirm given the poor fossil record of Myxiniformes. An anal fin could be a plesiomorphic characteristic of vertebrates if the ventral fin of *Myxinikela* is resolved as homologous to the anal fin of anaspids and gnathostomes. However, the distribution of the anal fin is variable among vertebrates [11]. The differentiation of an anal fin from a ventral fin fold might have occurred early in vertebrate history since anal fins supported by endoskeletal elements have been identified in fossil lampreys (e.g., *Hardistiella* and possibly *Mayomyzon*), despite its absence in extant forms (e.g, *Petromyzon, Lampreta*). In these, an anal ‘crest’ made of a skin fold devoid of fin rays occurs, but an anal fin may develop in certain atavistic specimens [54, 58]. There is no evidence of an anal fin in heterostracans [12] or arandaspids [59] so its absence can be considered a derived loss from the primitive condition of fossil lampreys. Janvier (2007; [54]) suggested that the ‘horizontal caudal lobe’ of osteostracans might be a modified anal fin. Galeaspids and pituriaspids appear to lack an anal fin [60, 61] but uncertainty is due to the poor preservation of their postcranial skeleton. In gnathostomes, the anal fin is not preserved usually in ‘placoderms’, probably due to its reduced size and the cartilaginous nature of the radials, but has been illustrated in some cases (e.g., arthrodires like *Africanaspis* and *Dunkleosteus*) [62, 63]). The anal fin is primitively present in all crown gnathostomes but may be absent in certain chondrichthyans [9, 11]. Living and fossil actinopterygians (with only a few exceptions in Osteoglossiformes, Anguilliformes, Lampridiformes, Siluriformes, and Syngnathiformes) and most piscine sarcopterygians (with exception in derived dipnoans) possess an anal fin. Among ‘elpistostegalians’, *Elpistostege* is known to retain an anal fin [15], which is definitely lost in tetrapods, while *Tiktaalik* might have lost the anal fin.

In the case of dorsal fins, paleontological and developmental evidence reveal that it is not constrained in its anterior extent and position, as opposed to the anal fin (which cannot extend anteriorly to the position of the anus), resulting in a variable occurrence of long-based and short-based dorsal fins in the earliest vertebrates [9]. *Pikaia* and Myllokunmingiida are the first and oldest early vertebrates in which a separate dorsal fin occurs. Lampreys display elongate dorsal fins that are not supported by radials. Extant lampreys display two dorsal fins, separated by a gap but the ‘posterior dorsal fin’ is now assumed to corresponds to the anterior extension of the caudal epichordal lobe seen in fossil lampreys (e.g., *Mesomyzon*; [64]). Fossil lampreys (e.g., *Hardistiella* and *Mayomyzon*) show no gap separating the anterior and posterior dorsal fins as in extant lampreys, suggesting that the double dorsal fins of lampreys and gnathosthomes is a convergent feature [54]. Duplication of short-based dorsal fins is also a common case in the evolution of vertebrates [9]. Several osteostracans have two dorsal fins (e.g., *Ateleaspis*, *Aceraspis*, and *Hirella*), but the anterior one lacks a fin web and resembles more a scale- covered hump than a proper fin [54]. On the other hand, the second dorsal fin of osteostracans clearly displays a fin web made of small scales arranged in a lepidotrichial pattern overlying numerous delicate radials. However, it is still debated whether this fin should be considered homologous to the posterior dorsal fin of gnathostomes or to the anterior part of the epichordal lobe of the caudal fin of lampreys, anaspids, and thelodonts. Among gnathostomes, ‘placoderms’ can either display single (e.g., antiarchs, stensionellids, rhenanids, and arthrodires) or double (e.g., ptyctodontids) dorsal fins [9]. Nevertheless, the plesiomorphic condition for crown gnathostomes, including chondrichthyans, ‘acanthodians’, and osteichthyans, is the occurrence of two dorsal fins supported by radials, fin rays, and sometimes associated spines [65–67]. Actinopterygians loss the anterior dorsal fin (from *Cheirolepis* onwards) but may regain a second dorsal fin either spinous (e.g., acanthomorphs) or adipous (e.g., euteleosts) [9]. Many sarcopterygians primitively retain two dorsal fins, with the exception of the derived loss of the dorsal fins in post-Devonian dipnoans and ‘elpistostegalians’ + tetrapods.

### Comparison of the median fin developmental patterning in osteichthyans

Fin and fin ray development have been well studied by developmental biologists since the middle of the 19^th^ century (e.g., [68–70]) and numerous studies have dealt with the morphological and molecular features of fin ray development and regeneration (see a review in [7] and references therein). Indeed, fin rays are a good tool to better understand vertebrate ontogenetic development [17] and the connections between gene expression (during normal development, regeneration, and mutagenesis) and morphological and structural variation of anatomical traits.

Fin and fin ray development has been thoroughly surveyed in osteichthyans through the zebrafish *Danio rerio* [7]. Other studies on fin anatomy and development have been performed mostly in extant actinopterygians such as *Salmo* [26], *Medaka* [71], *Tilapia* [72], *Amia* [73], *Polyodon* [6, 74], and *Acipenser* [74], but also in sarcopterygians such as the dipnoans *Neoceratodus* [75, 76], *Protopterus* [25], and *Lepidosiren* [77], and the coelacanth *Latimeria* [78, 79]. Our new data on extant (*Oncorhynchus*) and extinct (*Elonichthys, Miguashaia, Rhabdoderma, Dipterus, Quebecius,* and *Eusthenopteron*) osteichthyans allow us to accurately depict similarities in the developmental patterning of the median fins of bony fishes dealing with the morphological and temporary characteristics of appearance, chondrification, and ossification of endoskeletal elements (proximal and distal radials) and dermal fin rays (actinotrichia and lepidotrichia).

#### Apparition, chondrification, and ossification of radials and lepidotrichia

In extant actinopterygians (e.g., *Oncorhynchus*, *Salvelinus*, *Danio*) ([8,37,80]; this study) all developmental sequences (apparition, chondrification, and ossification) of the endoskeleton and exoskeleton for the dorsal and anal fins are bidirectional. This bidirectional pattern is corroborated by (1) common initiation sites for most corresponding events, (2) significant correlations between developmental sequences in both fins (dorsal and anal), and between the endoskeleton and exoskeleton (proximal/distal radials and lepidotrichia), and (3) a certain degree of simultaneity between sequences of apparition/chondrification (radials and lepidotrichia) and for the ossification of lepidotrichia. Despite little discrepancies among the initiation sites for apparition/chondrification/ossification of radials seen in *Oncorhynchus*, and between apparition/ossification of lepidotrichia, all these events can be confidently considered to be initiated from a unique initiation site. Unfortunately, the ossification patterns of many fossil osteichthyans are difficult to infer due to the preservation biases associated with the nature of fossilisation. However, in certain exceptional cases, ossification patterns can be tentatively reconstructed, such as in the coelacanth *Rhabdoderma* in which a bidirectional sequence occurred for the lepidotrichia Table 3, Fig. 8), whereas in the ‘osteolepiform’ *Eusthenopteron* ossification of the distal radials proceeded postero-anteriorly [17, 81] (Table 3); however, the narrowness of the fin and the reduced number of radials in *Eusthenopteron* might explain why the pattern is unidirectional.

In the case of lepidotrichia, fossil evidence confirms that dermal fin rays always ossify relatively early during ontogeny and before the endoskeletal radials ([82]; this study), a condition identical to that of *Oncorhynchus* and other extant osteichthyans. However, as for the radials, it is not easy to infer an ossification sequence for the lepidotrichia in immature fossil specimens.

#### Segmentation of lepidotrichia

Lepidotrichial growth is achieved by successive addition of distal segments at the extremity of the forming lepidotrichia [28]. The process of segmentation is congruent between the dorsal and anal fins in *Oncorhynchus* in terms of: (1) similar initiation sites, (2) common bidirectional sequences of segmentation, (3) simultaneity, and (4) significantly correlated sequences. The correlations among the lepidotrichia apparition, segmentation, and ossification sequences within both fins suggest that all three sequences are initiated from the same site and proceed in a bidirectional sequence. This congruence also matches the apparition/chondrification/ossification patterns described in radials.

In fossil taxa, an important aspect concerns the identification of the initiation site of segmentation. Considering the observation made in *Oncorhynchus* (i.e. longer lepidotrichia are the ones for which segmentation started earlier), the location of the longest lepidotrichia in the dorsal and anal fins can be confidently identified as the initiation site of segmentation in fossil specimens. This scenario has been proposed in all extinct osteichthyans surveyed (i.e., *Elonichthys*, *Miguashaia*, *Rhabdoderma*, *Quebecius, Dipterus,* and *Eusthenopteron*) (Table 3).

These similarities in patterning imply that sequences of segmentation may as well be generally used as proxies for sequences of ossification in osteichthyans. Considering the impracticality to observe an ossification sequence for lepidotrichia in fossil specimens, segmentation patterns are essential for comparisons between living and extinct taxa.

#### Bifurcation of lepidotrichia

Bifurcations are the results of the distal branching of an individual lepidotrichial segment after the intersegmental joint [38]. As for segmentation, bifurcation patterns can be compared in extant and extinct taxa. However, the bifurcation pattern is not clear in *Oncorhynchus*: (1) the sequences of bifurcation of both fins are bidirectional but not correlated, (2) the initiation site for bifurcation is posterior to the other initiation sites (apparition, ossification, segmentation) in the dorsal fin but similar in the anal fin, and (3) the initiation site corresponds to the lepidotrichia with the most proximal bifurcations in the dorsal fin but not in the anal fin. These results might be artefactual owing to small sample size and inter-individual variation. However, despite these potential biases, bifurcation sequences are likely similar between fins.

The bifurcation patterns found in both fins of extinct sarcopterygians (e.g., *Miguashaia*, *Quebecius*, *Dipterus*, *Eusthenopteron*) corroborate the observations made on the dorsal fin of *Oncorhynchus*: (1) an initiation site for bifurcation located posteriorly to the initiation site for segmentation, (2) concordance between the initiation site and the lepidotrichia with the most proximal bifurcations, and (3) a bidirectional sequence of bifurcation. These similarities between extant and extinct osteichthyans suggest a shared pattern of bifurcation in which bifurcation is initiated in a different position than the other events for the lepidotrichia.

### Median fin modularity and evolution in osteichthyans

Modularity is a fundamental property of organisms playing an important role in their evolution [83–85]. Anatomical modules refer to an internal organization of anatomical structures into distinct units, or modules, which develop and vary in quasi-autonomy, but within which the constituents interact and vary together [83,85,86]. This quasi-autonomy among modules allows three main evolutionary processes: dissociation, duplication/divergence, and co-option [83]. Different categories of modularity, and modules, have been defined over the past few decades. Among these categories, Zelditch & Goswami [85] emphasized the intricate relationship between developmental and functional modularity. Developmental modules are often represented as networks depicting their physical location, spatial extent and genetic specification, while functional modules are represented by anatomical elements integrated as structural components of a functional (or physiological) system [85]. However, developmental and functional modules have not been investigated methodologically as thoroughly as variational and evolutionary modules (see [85] for an exhaustive critical review of the methods). Thus, the identification of developmental and functional modules is frequently only suggested without being tested. Herein, we have proposed a method to compare similarities among sequences of developmental events helping to assess developmental modules.

Building on the original idea of Mabee et al. [16] of comparing sequence of formation (e.g., chondrification and ossification; e.g., in [87]), few studies have proposed to quantify the phenotypic patterning using correlation. Previous examples include correlation of relative sequence of events [88] and correlation of neural branching patterns in the skull [89]. Patterning is closely associated to the concept of developmental modularity [83, 89]. Thus, methodologically one has to compare similar patterning to infer developmental modules.

In our study, we are comparing our results that are methodologically constrained with previous developmental modules proposed by Mabee et al. [16] for actinopterygians: (1) the Endoskeleton and Exoskeleton Module (EEM) and (2) the Dorsal and Anal Fin Patterning Module (DAFPM). We will discuss the evidences supporting the occurrence of these modules in the median fins of the surveyed taxa, in osteichthyans, and in vertebrates as a whole.

#### Endoskeleton and Exoskeleton Module

The median and paired fins of osteichthyans are constituted of two skeletons, or modules, formed by distinct developmental processes: (1) the endoskeleton and (2) the exoskeleton [90–92]. In extant vertebrates, and more particularly in gnathostomes, the Endoskeleton and Exoskeleton Module (EEM) explains the similarities in the direction of development of the endoskeleton from the paired and median fins (fin radials and girdles) and the exoskeleton (fin rays). The EEM is thus composed of two interacting submodules (SM): (1) the endoskeleton submodule (EnSM) and (2) the exoskeleton submodule (ExSM). The EnSM as a whole probably originated at the base of the Gnathostomata with the evolution of pelvic fins in ‘placoderms’, homologous to those of osteichthyans [54, 93], while the ExSM is related to the origin of lepidotrichia in osteichthyans [24, 94] (Fig 12).

**Fig 12.**
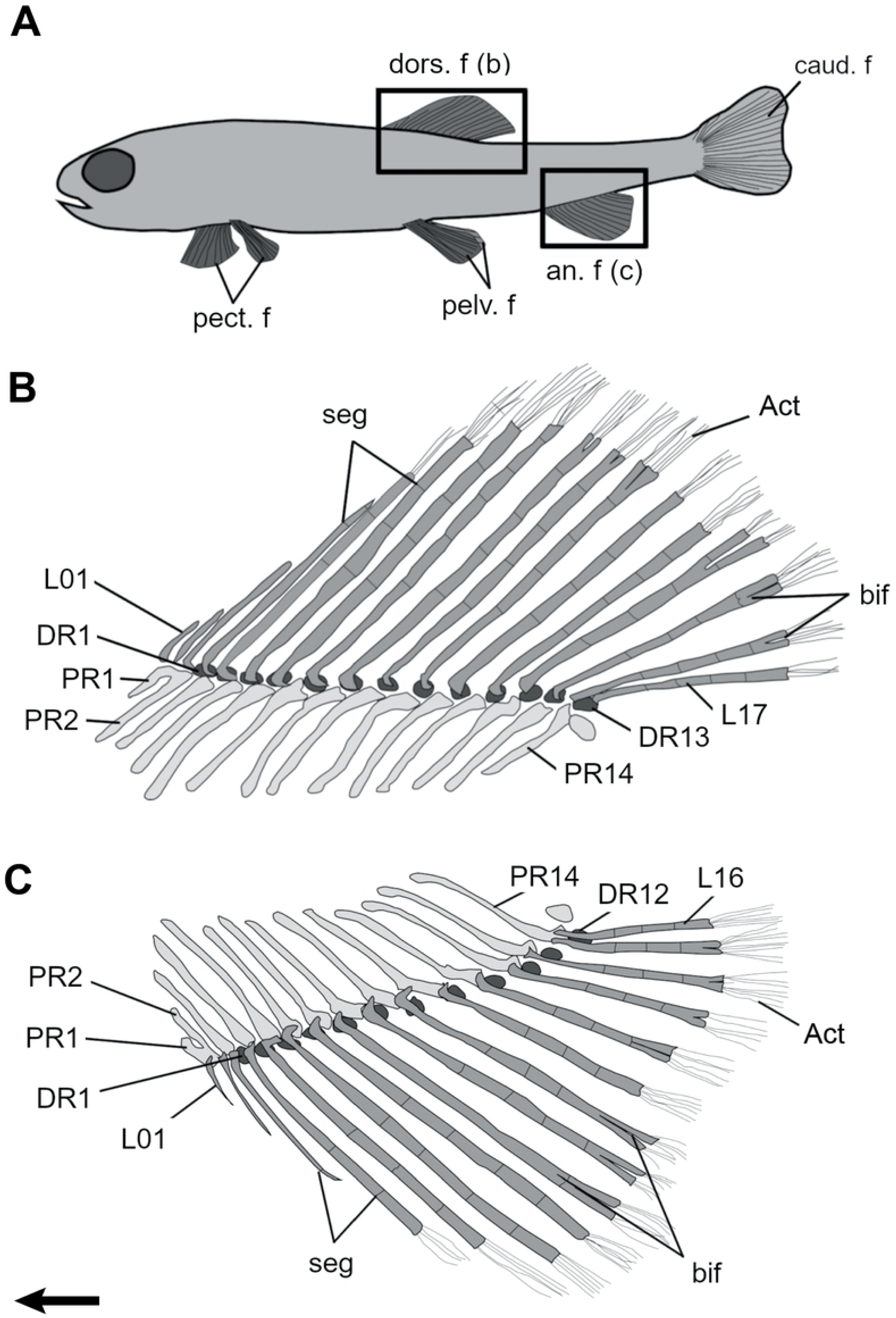
Vertebrate phylogeny illustrating the evolution of fin related characters and median fin modules. A, Interrelationships of the main groups of vertebrates and distribution of fin characters. Taxa in bold have living representatives. Fins have been plotted in the tree according to their definition as membranous outgrowths of the body walls internally supported by endoskeletal (e.g., radial bones) or exoskeletal (e.g., fin rays) elements and based on consensual hypotheses of homology (see main text for a discussion on the distribution of fins in the selected taxa). Preferred tree topology combined from [95–97] for non-gnathostomes, [98, 99] for actinopterygians, and [15, 100] for sarcopterygians; B, Distribution of median fin modules (DAFPM and EEM) in the studied species. DAFPM, dorsal and anal fin patterning module; EEM, endoskeleton exoskeleton module; LPM, lepidotrichia patterning module.

The endoskeletal components of a vertebrate fin includes series of radial bones, which support both paired (pectoral and pelvic) and median (dorsal, anal and caudal) fins. Endoskeletal radials have been proposed to be present in the median fins of the earliest vertebrates (e.g., myllokunmingiids like *Haikouichthys*) [16, 101]. However, these supposedly cartilaginous structures have been reinterpreted as either epidermal folds or collagenous structures [11,54,102] and the presence of true radials is thus now restricted to crown vertebrates as present in the caudal fin of hagfishes, lampreys, and even conodonts [11,54,103]. Radial elements (either osseous or cartilaginous) have been identified in the caudal fin of many other subsequent groups of vertebrates (e.g., heterostracans, anaspids, osteostracans and gnathostomes) [52], but it is not always clear whether the other median fins (i.e., dorsal and anal) were also supported by radials. The ribbon-shaped anteroventral paired fins of certain ‘agnathans’ (e.g., *Euphanerops*, anaspids) present numerous thin, parallel radials, lacking any fin support or girdle [55], whereas in osteostracans, possibly pituriaspids, and gnathostomes a few large radials articulate with a massive girdle forming stout, paddle-shaped paired fins [11, 54].

Dermal fin rays are absent in cyclostomes (hagfishes and lampreys), but they are known in non-osteichthyans (e.g., in ‘agnathans’ like *Euphanerops* and in gnathostomes like ‘acanthodians’ and chondrichthyans) [55,63,104,105]. However, the structure and histological nature of the fin rays in early vertebrates is difficult to decipher. In *Euphanerops*, the paired anteroventral fins are composed of ‘fin rays’ made of stacked chondrocytes, which articulate with the cartilaginous radials [55]. In anaspids (e.g., *Birkenia*), the anal and caudal fin are covered with small scales and in the epichordal lobe of the caudal fin the scales are arranged in rows, recalling the jointed structure of the lepidotrichia in osteichthyans [52]. The small median fins of thelodonts (e.g, *Phlebolepis*) are also covered by minute scales, closely stacked together and sometimes forming a fin web [106], similar to the configuration of the caudal fin of the arandaspid *Sacabambaspis* [58]. In osteostracans (e.g., *Escuminaspis*), the second dorsal fin is supported by numerous narrow radials and covered by small scales arranged in rows [52, 107]. Among gnathostomes, chondrichthyans possess ceratotrichia [104,108–111], large fibrous fin rays but homologous in all senses to the actinotrichia of osteichthyans [19,78,112,113].

Ceratotrichia have also been identified in ‘placoderms’ [e.g., *Bothriolepis* [114–117], dunkleosteids [118], stensionellids [115]. ‘Acanthodians’ possess dermal fin rays with an ossified proximal portion and a distal, non-ossified portion, which, according to Géraudie and Meunier [78] correspond to large fibrous rays, similar to ceratotrichia. Wide ceratotrichia may thus represent the primitive condition from which the slender actinotrichia evolved in osteichthyans. Completely ossified fin rays (lepidotrichia) are solely present in osteichthyans.

The exoskeleton submodule (ExSM) might thus have been present in the last common ancestor of chondrichthyans and osteichthyans, since partially ossified fin rays (i.e., potential lepidotrichia) may be present in the caudal fin of certain ‘acanthodians’ [119, 120], currently considered as stem chondrichthyans [65, 121,122]. On the other hand, this module is lost in the limbs of tetrapods [91, 123], in which dermal fin rays are absent from the paired fins but still retained in the caudal fin of Devonian forms (e.g., *Acanthostega, Ichthyostega*) [124, 125]. This pattern of lepidotrichial reduction at the transition between fishes and tetrapods has been documented in *Tiktaalik* by looking at the dorso-ventral asymmetry of hemirays [126]. The origin of the EEM most likely coincides with the common presence of both submodules (EnSM and ExSM) in crown gnathostomes (Fig. 12). Interactions between these two modules resulted in the morphological disparity in terms of relative size, shape, and position of the paired and median fins.

Based on the sequence of nine events surveyed during *Oncorhynchus* fin development, the patterning of the EEM in living actinopterygians is composed of five events, all starting from the same initiation site and following the same direction: (1) apparition/chondrification of radials, (2) apparition of lepidotrichia, (3) segmentation of lepidotrichia, (4) ossification of lepidotrichia, and (5) ossification of proximal radials. The ossification of distal radials is probably the sixth step, but it was not observed in our juvenile specimens. Based on our observations, the EEM can thus be confidently considered plesiomorphic at least in actinopterygians, since it was not possible to confirm its presence in fossil sarcopterygians.

#### Dorsal and Anal Fin Patterning Module

The anatomical composition, disparity, phylogenetic distribution, and modularity of fins in all orders of extinct and living fishes have been surveyed by Larouche et al. [9, 11]. Larouche et al. [9] recognized that the dorsal and anal fins form an evolutionary module nested within a median fin module. A Dorsal and Anal Fin Module (DAFM) most likely evolved early in stem-gnathostomes (Fig. 12). Larouche et al. [11] suggested that fin modules, including DAFM, re-expressed within the topographic boundaries of fin-forming morphogenetic fields.

Mabee et al. [16] described the DAFPM based on a similar direction of differentiation (i.e., apparition of the skeletal elements) for the dorsal and anal fins of actinopterygians. However, our data allow to expand Mabee et al.’s original description of the DAFPM to also include the patternings of chondrification, ossification, segmentation, and bifurcation (Tables 2 and 3; Fig 3). Further evidences of the DAFPM are given on extant actinopterygians by similar coordination of phenotypic plastic response between both fins [37] and similar pattern of correlated shape between both fins [127]. Mabee et al. [16] considered the bidirectional direction of development as plesiomorphic at least for teleosts. Our data show that the sequences of all events are effectively bidirectional in *Oncorhynchus* and most fossil osteichthyans (Table 3) thus suggesting that developmental bidirectionality is most likely plesiomorphic for osteichthyans. This general bidirectional development may differ in some sarcopterygians (e.g., radials in *Eusthenopteron*), most likely owing to a reduction of size in both fins, but does not necessarily compromise the presence of the primitively shared DAFPM among osteichthyans.

The Dorsal and Anal Fin Patterning Module (DAFPM) is the developmental module corresponding to the anatomical DAFM of Larouche et al. [9]. In addition, Mabee et al. [16] suggested that the DAFPM originated from a Dorsal and Anal Fin Positioning Module (DAFPoM), one in which the antero-posterior position of the dorsal and anal fins are correlated. Most likely the basal condition of the DAFPoM in actinopterygians is a condition in which these median fins occupy a symmetrical position [16]; this condition is observable in Devonian actinopterygians (e.g., *Dialipina, Pickeringius, Howqualepis, Mimipiscis* [128–130]), while the anal fin is located slightly anterior to the dorsal fin in *Cheirolepis* [131] or the opposite (e.g., *Gogosardina, Moythomasia, Limnomis* [132–134]). Mabee et al. [16] mentioned that there was a high level of dissociability of the positioning module from the patterning module among actinopterygians. Outside actinopterygians, the generalized condition in actinistians (*Miguashaia*; [22]), dipnomorphs (e.g., *Holoptychius, Quebecius, Glyptolepis, Uranolophus*; [48, 49]) and tetrapodomorphs (e.g., *Cabonnichthys, Eusthenopteron, Gyroptychius, Heddleichthys*; [50,135,136]) also corresponds to the symmetrical positioning. Not only the position of the dorsal and anal fins is symmetrical, but the shape, size and number of endoskeletal elements correspond to a mirror image. The DAFPM of osteichthyans implies that the skeletal elements (either of endochondral origin such as the radials or dermal origin such as the lepidotrichia) of both median fins (anal and dorsal) differentiate in the same direction and thus share common developmental properties [16].

The DAFPM is confirmed by our data in actinopterygians (*Oncorhynchus*, *Elonichthys*), as well as in sarcopterygians like coelacanths (*Miguashaia*, *Rhabdoderma*), porolepiforms (*Quebecius*), and ‘osteolepiforms’ (*Eusthenopteron*) (Tables 2 and 3). The DAFPM can also be recognised in early lungfishes (e.g., *Dipterus, Barwickia*) based on morphological similarities between these fins, particularly with respect to the oar-shaped fin support supporting the distal radials [137]. However, due to the heterogeneity of fin morphologies in dipnoans it is difficult to confidently reconstruct the evolution of the DAFPM across lungfishes [17]. Consequently, a dissociation of the DAFPM is inferred during dipnoan evolution. Multiple dissociations have also occurred in actinopterygians every time either one of the dorsal or anal fin is absent or fused to the caudal fin (e.g., Osteoglossiformes, Anguiliiformes, Siluriformes, Lampridiformes, Sygnathiformes) [9]. Among other sarcopterygians non-surveyed in our study, the DAFPM can be certainly inferred in onychodonts (e.g., [138]) and rhizodonts (e.g., [139]) based on the size and shape similarities between both median fins. The DAFPM is lost definitely at the basis of the clade including the ‘elpistostegalians’ and tetrapods with the loss of the dorsal fins (Fig 12).

#### Lepidotrichial Patterning Module

Molecular mechanisms and grafting experiments in zebrafish suggest that a lepidotrichium can be grafted to a new location and grows quite normally by the addition of new segments [140, 141]. However, interactions among lepidotrichia are necessary to achieve the original morphology because the position of segmentation and bifurcation are dependant of the position of the lepidotrichia within fins [140, 141]. This suggests a smaller modular unit within the exoskeleton module (ExM), the lepidotrichium itself. The hemi- lepidotrichium (or hemiray) could be the smallest unit of regeneration (and development) in the hierarchical modular organization of fins [142], because developmental interactions are recognized to control coordination of segmentation and bifurcation between hemirays [141]. Segmentation and bifurcation patterning similarities found in this study may be indicative of molecular mechanism conservatism in osteichthyans.

Lepidotrichia patterning in *Oncorhynchus* includes: (1) apparition, (2) segmentation, (3) ossification, and (4) bifurcation. Data from fossil specimens agree with this sequence. Therefore, the patterning of the lepidotrichia appears conserved in osteichthyans and a “Lepidotrichia Patterning Module” (LPM) may be generalized in all the fins of all osteichthyans.

The molecular basis behind the establishment of segmentation has been explored by using regenerative experiments in zebrafish. Part of the molecular machinery involved in segmentation is recruited for lepidotrichia bifurcation [18, 143]. Molecular transcripts involved in segmentation are: (1) *evx1* acts as an on/off switch defining the putative boundaries between two successive segments [28], mutant zebrafish for this gene grow normal lepidotrichia, but joint formation between successive segments is impaired [38]; (2) three genes of the Sonic Hedge Hog signalling pathway (i.e., *shh*, *bmp2, ptc1*) are thought to be involved in the patterning of the lateral limits of segments [18]; (3) *hoxa13b* may participate in the elaboration of the next segment [144]; and (4) *cx43*, a particular gene allowing intercellular communication involved in joint position [145].

Segmentation and bifurcation processes might be independently regulated. In *evx1* zebrafish mutants, bifurcation occurs normally while joint formation is down-regulated [38]. At least two genes are expressed during the bifurcation process: (1) prior to bifurcation, *shh* is expressed centrally where a new segment is forming, then laterally in the presumptive twin-segments [18, 143], and (2) *bmp2* (required for bone synthesis in the central region of the segment) is restricted in the two lateral domains copying *shh* prior to branching [18]. Fossil evidence shows that bifurcation likely originated at the base of osteichthyans. One notable exception is the puzzling *Dialipina salgueiroensis,* which lacks bifurcation (RC, pers. obs.); *Dialipina* is either considered a stem actinopterygian [146] or a stem osteichthyan [24]. The absence of bifurcation is highly homoplastic in living [32, 147] and fossil actinopterygians as well as in some sarcopterygians (e.g., it constitutes a common derived feature of post-Devonian coelacanths) [148]. Because bifurcation is the last event of the lepidotrichia developmental sequence, it might be more susceptible to be affected by epigenetic phenomena [37].

## Conclusions

Our analysis of median fin development in *Oncorhynchus* has allowed the quantification and validation of two median fin modules in a living actinopterygian: the Dorsal and Anal Fin Patterning Module (DAFPM) and the Endoskeleton and Exoskeleton Module (EEM). Comparison with other extinct osteichthyans, comprising both actinopterygians and sarcopterygians (coelacanths, dipnoans, porolepiforms and ‘osteolepiforms’) has corroborated the data on extant taxa, but highlighted the difficulties of confidently identifying developmental sequences based on fossil specimens. The DAFPM and EEM modules incorporate all the events associated with fin patterning including the sequences of segmentation and bifurcation of the lepidotrichia that are crucial for comparisons and inferences of developmental sequences in fossil osteichthyans.

We suggest that: (1) the EEM includes the apparition, segmentation, and ossification sequences, and, based on our results, is plesiomorphic at least for actinopterygians; (2) the DAFPM includes the apparition, chondrification, and ossification sequences plus the segmentation and bifurcation sequences, and is plesiomorphic for osteichthyans with multiple dissociations along osteichthyan phylogeny. Additionally, the recurrence of the developmental pattern of the lepidotrichia in living and fossil osteichthyans suggests an additional developmental module within fins, the Lepidotrichia Patterning Module (LPM), where the constitutive units, the hemirays, have a synchronous and similar development. The median fins of osteichthyans have thus been shown to be important representatives for the study of modularity across the evolution of vertebrates.

## Supporting information

**S1 Table. Fossil specimens examined given in size order.** For each species, the size series is given with SL or TL (mm) and specimen number. Only *D. valenciennesi* is not represented by immature specimens.

S1 File. Coding of the serial skeletal elements, i.e. the proximal radials (PR), the distal radials (DR) and the lepidotrichia (L), during nine out of eleven developmental events of the developmental sequence of the dorsal and anal fins of *Oncorhynchus mykiss*. For each specimen, the size is given in SL (mm) and the skeletal elements are coded with 0, 1 or NA. The definitions of these codes are different for each developmental event, they are provided.

## Acknowledgments

For the loan of specimens, we thank J. Maisey, I. Rutzky and J. Galkin (AMNH), L. Grande, B. Simpson and E. Zeiger (FMNH), J. Kerr, N. Parent and J. Willett (MHNM), A. Lévesque (ULQ), and J. Gauthier and M.A. Turner (YPM). S. Cumbaa (NMC), Z. Johanson (NHM), and K. Mickle (KU) provided specimen photographies. I. Béchard digitized the drawings. I. Béchard, L. Fischer-Rousseau, Z. Johanson, and C. Riley provided constructive comments on earlier versions of this manuscript. B. Vincent (UQAR) provided statistical and programming advices. P. Janvier (MNHN) and M. Marí-Beffa (UMA) are warmly thanked for their insightful comments on fin morphogenesis, fin ray structure and fin diversity.

